# Distinct 2-phenyl-imidazo[1, 2α] pyridine derivatives drive ER degradation and selectively impair proliferation of ER^+^ breast cancer cells via the aryl hydrocarbon receptor

**DOI:** 10.1101/2025.07.28.666967

**Authors:** Katrin Koellisch, Christine Blattner, Stefano Motta, Janine Wesslowski, Melanie Rothley, Simone Büchel, Savannah Sirounian, Julia Müller, Marina Grimaldi, Jutta Stober, Zoe Wammetsberger, Mengwu Pan, René Houtman, Christoph W. Grathwol, Ilenia Segatto, Lo-Wei Lin, Laki Buluwela, Siva Kumar Kolluri, Simak Ali, Nicole Jung, Barbara Belletti, Laura Bonati, Patrick Balaguer, William Bourguet, Stefan Bräse, Gary Davidson, Andrew C. B. Cato

## Abstract

X15695 is a 2-phenyl-imidazo[1, 2α] pyridine derivative identified as an orally active, selective oestrogen receptor (ER) degrader that inhibits the proliferation of ER^+^ breast cancer cells. Here, we show that X15695 is an aryl hydrocarbon receptor (AHR) ligand that stabilises the AHR more efficiently than its classical ligand, indirubin. X15695 enables AHR to form a complex with the ER, promoting its proteasomal degradation. In the presence of oestradiol, X15695 outperforms the standard of care drug fulvestrant in suppressing the growth of ER^+^ breast cancer cells, either expressing the wild-type or clinically relevant ER mutant forms (Y537S and D538G), and of patient-derived xenograft organoids established from ER^+^ tumours. Using computational techniques, we discovered that a low pKa value resulting from electron-withdrawing substituents in the 2-phenyl-imidazo[1, 2α] pyridine compounds is a key feature that identify them as potent AHR ligands, leading to the potential discovery of additional derivatives for future therapeutic development.

## Introduction

Breast cancer is one of the most frequently diagnosed cancers and the leading cause of cancer death in women worldwide ^1^. About 70% of all breast cancers express the hormone receptors, oestrogen (ER) and/or progesterone (PR) receptors, and are negative for human epidermal growth factor receptor 2 (HER2) and rely on hormone signalling for growth ^2^. Current therapies that target this axis in the adjuvant setting include inhibitors of oestrogen production such as luteinizing hormone-releasing hormone (LHRH) agonists, aromatase inhibitors (AIs), or selective oestrogen receptor modulators (SERMs) such as tamoxifen and selective oestrogen receptor degraders (SERDs) such as fulvestrant ^3^.

While these endocrine therapies have significantly improved the outcome of hormone receptor-positive breast cancer, both *de novo* and acquired endocrine resistance remain a major challenge ^4^. Acquired resistance is frequently caused by increased mutations in the ligand binding domain of the gene coding for oestrogen receptor alpha (*ESR1*). Such mutations are rare in primary or treatment-naive primary tumours but frequently arise under therapies that target oestrogen signalling pathways ^5,6^. The *ESR1* mutations drive oestrogen-independent constitutive activation of ER and oestrogen-independent growth ^7^ and are detected in nearly 30% of all ER^+^ metastatic patients. Of these, more than a half are accounted for by the variant Y537S (21%) and D538G (33%) ^8^. Among all *ESR1* mutations, the Y537S mutation, in particular, is associated with a higher degree of resistance to most endocrine therapies ^7–10^.

Fulvestrant, the first clinically approved SERD showed only modest inhibition of mutant ERα, compared to wild-type ERα ^3^. Besides, fulvestrant is limited in its clinical use by its intramuscular formulation and once-a-month injection ^11^. A new class of inhibitors with superior bioavailability and ER-degrading potential (oral SERDs) such as elacestrant, giredestrant, amcenestrant and camizestrant have therefore been developed ^11,12^. Elacestrant has been approved in postmenopausal women or adult men with ER^+^/HER2^-^, *ESR1* mutated advanced or metastatic breast cancer following prior endocrine therapy ^13^. Camizestrant might become the second oral SERD to be approved in metastatic disease following the positive results it received in the SERENA-2 study where it showed a significant benefit in progression-free survival (PSF) versus fulvestrant ^14^. Unfortunately, in the AMEERA-3 trial that compared amcenestrant with endocrine treatment of physiciańs choice in second or third-line metastatic setting, an improvement in PFS was not demonstrated ^15^ and its development was therefore discontinued. A more positive outcome may come from the acelERA breast cancer study comparing giredestrant with physiciańs choice of endocrine monotherapy. Although no statistical significance was reached in the primary investigator-assessed PFS ^16^, a trend toward favourable benefit among patients with *ESR1*-mutated tumours was observed, supporting the continued investigation of giredestrant in other studies. Other SERDs are still in earlier stages of development ^12^ but on the whole, the benefits of oral SERDs in the premenopausal population have not yet been demonstrated and there is therefore the need for future development of oral SERDs for these indications.

We have recently described 2-phenyl-imidazo[1, 2α] pyridine derivatives as potent inhibitors of breast and prostate cancer cell growth in preclinical studies. The prototype compound X15695 was shown to inhibit MCF-7 breast cancer xenografts and to degrade ERα in *in vitro* cell culture experiments and in *in vivo* studies when administered orally and it was therefore classified as an oral SERD ^17^. Apart from its ability to degrade ERα, X15695 was shown to activate p53 and to promote cell cycle block and induce apoptosis. However, the mode of action of this compound class is unclear and its primary target has not been identified. In this study, we have identified X15695 and additional 2-phenyl-imidazo[1, 2α] pyridine derivatives as ligands of the aryl hydrocarbon receptor (AHR) and shown that X15695 strongly interacted with AHR, enhanced its transactivation function, formed a complex with ERα and destabilised ERα via the proteasomal pathway. In the presence of oestradiol, X15695 outperformed fulvestrant in inhibiting the proliferation of MCF-7 cells expressing the clinically relevant ER mutations that confer resistance to current endocrine therapies. Importantly, these favourable profiles of X15695 were also confirmed in preclinical model, using organoids from patient-derived xenograft (PDX) established from an ER^+^ breast cancer patient. We have therefore identified a unique property of a SERD that can be harnessed in future therapeutic developments.

## Material and Methods

### Drugs and Chemicals

17 7-oestradiol (Sigma Aldrich, E8875; PubChem SID 24278426), fulvestrant (Sigma Aldrich, I4409; PubChem SID 329815373), indirubin (Sigma Aldrich, SML0280; PubChem SID 329825288), Dioxin (Campro Scientific), Lipofectamine 2000 (Thermo Fisher 11668027), Protein A-agarose beads (Thermo Fisher, 20333), Protein G-agarose beads (Thermo Fisher 11668027), CH-223191 (MedChemExpress, PubChem SID 329775161), Epoxomicin (UBPBio, Pub Chem SID 329799052). Synthesis of the 2-phenyl-imidazo[1, 2α] pyridine derivatives used in this study have been previously described ^17^.

### Cell lines

All cell lines except HAhLH cells were obtained from the American Type Culture Collection (ATCC): MCF-7 (RRID:CVCL_0031), T47D cells (RRID: CVCL_0553), HeLa (CCL-2, RRID:CVCL_0030). CRISPR ER knock-in cell lines (MCF-7 luc, MCF-7 luc D538G and MCF-7 luc Y537S) have been reported previously ^18^. Cells were all mycoplasma-free upon receipt from ATCC prior to 2015. The identities of the cells were confirmed by short tandem repeat profiling (BioSynthesis, Lewisville, TX and DSMZ Braunschweig, Germany). All cell lines are routinely confirmed to be *mycoplasma*-free, using the Venor®GeM Classic Mycoplasma Detection Kit for conventional PCR (Minerva Biolabs, 11-1250). MCF-7 were cultured in DMEM supplemented with 10% foetal bovine serum (FBS), 1% penicillin/streptomycin, T47D, MCF-7 luc, MCF-7 luc D538G and MCF-7 luc Y537S were cultured in RPMI 1640 medium supplemented with 10% foetal bovine serum (FBS), 1% penicillin/streptomycin. HAhLH cell line was cultured in DMEM-F12 supplemented with 5% foetal bovine serum (FBS), 1% penicillin/streptomycin, 0.25mg/ml hygromycin. All cells were maintained at 37 °C in an incubator with 5% CO_2_ and 90% humidity. Unless otherwise stated, for all experiments requiring hormone starvation, cells were cultured for 72 h in phenol red-free RPMI 1640 and DMEM media respectively, supplemented with 3% charcoal-stripped foetal calf serum (CCS).

### Generation of Patient Derived xenograft (PDX)-derived Organoids (PDxO) Ethical statement

Human specimens were collected from breast cancer patients undergoing surgery at the National Cancer Institute, Aviano, upon signing an informed consent form. The research was permitted by the Institutional Review Board of CRO Aviano (IRB-06-2017) and complied with all relevant ethical regulations, including the Declaration of Helsinki.

### Procedure

Organoids were generated from patient-derived xenograft (PDX) tumours, as previously described ^19^. PDX were derived from BCRO#28, an ER^+^ breast cancer sample collected from a treatment naïve 57 years-old patient. Briefly, tumours freshly explanted from mice were mechanically shredded, digested with collagenase (Merck) and filtered through a 100 μm strainer. Digested material (isolated cells and small cell clusters) was centrifuged and embedded in 3D-matrix drops (reduced growth factor Basement Membrane Extract, BME, R&D systems), then overlaid with breast cancer organoid medium ^19^. For 3D survival curves under treatment, organoids were embedded in BME drops and, after 24-48 hours, drugs were added. Fresh drug-containing medium was replaced at day 4. At day 7 of treatment, CellTiter-Glo® 3D Cell Viability Assay (Promega) was added and luminescent signal from viable organoids was recorded on Infinite M1000 Pro instrument (Tecan).

### Reporter cell line assay

The reporter HAhLH cell line was obtained by transfecting human HeLa cells with the dioxin-responsive gene XRE(TnGCGTG)3-tata-luciferase-Luc-hygromycin plasmid as previously described ^20^. For measuring AHR activity of chemicals, the cells were seeded in 96-well white opaque tissue culture plates at a density of 50,000 cells per well in 150 μl of culture medium. Test compounds were prepared at 4x concentration in the same medium, and 50 μl per well were added 24 h after seeding. Cells were incubated for 8 h in the presence of the compounds at 37 °C. At the end of incubation, the medium containing test compounds was replaced with test culture medium containing 0.3 mM luciferin and luminescence was measured in intact living cells for 2 s. Experiments were performed in quadruplicate for each ligand concentration. Each compound was tested in at least four independent experiments. Results were expressed as a percentage of maximal luciferase activity. Maximal luciferase activity (100%) was obtained in the presence of 10 nM dioxin. Effective concentration (ECs) for a given compound, EC_50_ is defined as the concentration inducing 50% of its maximal effect. The EC_50_ values were calculated including the adjustment for the basal activity of the cell line.

### CRISPR/Cas9 knock out

The human AHR locus was targeted within exon 1 to introduce a frame shift. Using the CRISPOR online tool ^21^ two 20 bp protospacer (805 5’-CCTACGCCAGTCGCAAGCGG-3’, 708 5’-CCAGCCTACACCGGGTTCCG-3’) followed on the 3’ end by a NGG PAM (protospacer adjacent motif) upstream and downstream of the AHR start codon were identified. DNA oligonucleotides were designed and annealed to generate short double-stranded DNA fragments containing the 20 bp protospacer and an additional guanine nucleotide at the 5’end was added to increase targeting efficiency ^22^ and 4 bp overhangs suitable for ligation into a BbsI restriction site. This short double-stranded DNA fragments were ligated into the BbsI restriction site of pSpCas9(BB)-2A-GFP (PX458) vector, a gift from Feng Zhang (#48138, Addgene). To knock out AHR, T47D cells cultured in 6-well plates were transfected with 2 µg pSpCas9(805)-2A-GFP to induce a frame shift or with 1 µg pSpCas9(805)-2A-GFP and 1µg pSpCas9(708)-2A-GFP to delete ATG using ScreenFectA (ScreenFect GmbH) according to the manufacturer’s 1-step protocol. MCF-7 cells were transfected using Lipofectamine™ 3000 (Thermo Fisher Scientific). 48 h after transfection, GFP-positive cells were sorted by FACS, and single cell clones were expanded for further characterization. After SDS-PAGE and Western blot screening of the FACS sorted clonal cell lines, genomic DNA from promising candidates was PCR amplified at the region of interest (F 5’-CCAGGCAGCTCACCTGTAC -3’; R 5’-CTCTAACCTAACCCATGCGGATATG -3’) and sent for sequencing (Microsynth AG, Balgach, Switzerland).

### Western blotting and Co-immunoprecipitation

For Western blotting, cells were washed with ice-cold PBS and lysed in NP40 lysis buffer (1% NP-40, 50 mM Tris-HCl, pH 8.0, 150 mM NaCl, 5 mM Ethylenediaminetetraacetic acid [EDTA], 1 mM PMSF) followed by five pulses of sonication at 50 amplitudes. Equal amounts of protein were separated by a 9% SDS-PAGE gel. Western blotting was carried out using standard protocols with the following antibodies: ERα (Santa Cruz Biotechnology, catalogue no. sc-8002, <u>RRID:AB</u>_627558), PCNA (Santa Cruz Biotechnology, catalogue no. sc-56, RRID:AB_628110), Goat anti-mouse Immunoglobulins/HRP (DAKO, catalogue no. P0447, RRID:AB_2617137).

For co-immunoprecipitation experiments, 1×10^6^ MCF-7 cells were seeded in 15 cm dishes and cultured for 1 day in minus phenol red medium supplemented with 3% CCS. Cells were treated with 10 nM oestradiol (E_2_), 1 μM X15695, E_2_ and E_2_ + X15695 or the equivalent volume of DMSO for 1 h. Cells were harvested, washed with phosphate buffered saline (PBS) and lysed using IGEPAL-CA-630 buffer (50 mM Tris HCl pH 7.4; 0.5 % IGEPAL-CA-630; 10 mM EDTA; 450 mM NaCl; 50 mM NaF; 1 % PMSF; 10 % Glycerol). Protein A agarose beads were mixed with protein G agarose beads (4:1) and incubated for 30 min at room temperature with anti-AHR or control IgG antibody. Cellular extracts from treated and untreated cells (1,900 μg) were incubated with antibody-preincubated protein A/G agarose beads at 4 °C for 2 h. The beads with immunoprecipitated proteins were collected at 4 °C by centrifugation at 1,000 g for 5 min. and washed 4 to 5 times with PBS. The immunoprecipitated proteins were eluted by heating in SDS sample buffer for 5 min at 95 °C and resolved on SDS-PAGE. Thereafter, they were subjected to Western blot analysis to detect ERα. Cellular extracts containing 30 μg protein from treated and untreated cells were used as the input control for the targeted proteins.

### Fluorescence polarisation assay

The PolarScreen™ Estrogen Receptor-Alpha Competitor Assay, Red (ThermoFisher, A15884) was used to determine if the X compounds bind to ERα. The assay was performed according to the manufacturer’s protocol. In brief, 34 nM ERα full-length and 1.4 nM Fluormone™ EL Red were incubated for 4 h at room temperature together with known concentrations of oestradiol, fulvestrant or the X compounds. Fluorescence polarisation was measured in a Tecan F200 reader with 535 nm excitation and 595 nm emission interference filters.

### Clonogenic assay

Cells were seeded at a density of 1-2×10^3^ cells/well in a 6-well plate and treated with increasing concentrations of the X compound and cultured for 14-21 days. Medium and compounds were exchanged every 7 days. Cells were fixed with methanol and the formation of colonies was visualized using 0.5% crystal violet (w/v) in 20% methanol (v/v). Plates were photographed and the area covered by colonies was calculated using the ColonyArea Plugin for ImageJ (RRID:SCR_003070) ^23^.

### Cell Proliferation Assay

For the proliferation assay, indicated cell lines were plated in a 24-well plate format at 1×10^4^ cells/well. Cells were cultured for 5 days and cell growth was determined by trypan blue exclusion using the direct cell count function on a Countess II (Life technologies).

### RNA sequencing and quantitative RT-PCR experiments

RNA sequencing (RNA-seq) has previously been carried out with MCF-7 and T47D cells treated with 10 nM 17-β-oestradiol (E_2_) and 1 µM X15695 and the results have been deposited at the GEO repository under accession number GSE218556. The RNA-seq datasets were analysed using a fold change of Log_2_(CPM) ≥ 1.0 or ≤ -1.0 and an adj. p value ≤ 0.05 to identify differentially expressed genes (DEGs) in response to X15695 in the presence or absence of E_2_. Gene Ontology (GO) analysis and Gene set enrichment analysis (RRID:SCR_006484) were performed using R package GSEABase (v1.58.0) and GSVA (v1.44.2) and the results were visualized by R package enrichplot (v1.16.1) ^24,25^. The DEGs enriched in the pathway were used to generate heatmaps with the heatmap2 package. Select target genes were analysed by RT-PCR experiments. PCR analysis was carried out using the SYBR Green GoTaq PCR mix (Promega, A6002) and βactin as a housekeeping gene. The sequence of all the primers used can be found in Supplementary Table 1.

### Thermal stability measurements (nano-DSF)

Thermal denaturation of the AHR complexes in the presence of compounds was analysed using a Tycho NT.6 device (NanoTemper Technologies). The protein sample was prepared as previously described ^26^ and diluted in the buffer containing 20 mM Bis-Tris–HCl pH 7.0, 50 mM NaCl, 10 mM KCl, 10 mM MgCl_2_, 20 mM Na_2_MoO_4_ and 2 mM BME to a final protein concentration of 2.5 μM. Compound solutions at 20 mM were prepared in DMSO and added to the sample to a concentration of 12.5 μM. DMSO alone was used as a reference at a concentration of 5% (v/v). Upon addition of the ligand, samples were incubated at room temperature for 10 min. Denaturation curves were recorded in a range from 35 to 95 °C with 1 °C/min steps and plotted as a first derivative of the fluorescence ratio at 330 nm/350 nm against temperature. All measurements were performed in triplicates.

### Nuclear receptor Activity Profiling (NAPing)

Preparation of assay mixes with ERα recombinant ligand binding domain (LBD) and full-length ERα in MCF-7 extracts and measurement of coregulator recruitment on the NAPing platform (PML, Oss, The Netherlands) was performed as described previously ^27,28^. In short, samples with ERα LBD-GST (Invitrogen #PV4543) or full-length protein were stimulated with -8.5 logM (∼EC_50_) of oestradiol (E_2_) and subsequently incubated with 1000-fold excess (-5.5 logM) of each test compound and incubated for 30 min at room temperature on the NAPing platform with 101 coregulator-derived binding NR-binding motifs. ERα binding was detected using ALEXA488-GST (Invitrogen #A-11131, for LBD) or ERα /Goat-anti-Rat-FITC (Santa Cruz sc-53490/Novus Bio NB7124) antibodies. Unbound ERα and antibodies were removed by washing. Residual ERα binding to each coregulator motifs was recorded on tiff images and quantified using an R-based analysis software package (PML, Oss, The Netherlands).

### Determination of the thermodynamic solubility

Thermodynamic solubility was determined using the classical shake-flask method. The solid compound (4 mg) was suspended in 1 ml HEPES buffer (0.1 M, pH 7.4) and constantly stirred at 21 °C for 24 h. Thereafter, the suspension was centrifuged at 13,000 rpm for 5 min (Centrifuge Mikro20, Andreas Hettich GmbH) and the clear supernatant was transferred to an HPLC vial (flat bottom, borosilicate glass 5.1, 1.5 ml, ND9 (Cat. No. 548-0028AP, VWR International) in combination with 9 mm natural rubber/TEF screw caps (Cat. No. 11722438, Fisher Scientific) and analysed using an Agilent Single Quadrupole LC/MSD iQ Mass-System equipped with a Kinetex 2.6 µm XB-C18 100 Å LC column (100 x 4.6 mm). The Absorption (254 nm) was measured and the Area Under the Curve evaluated (AUC). For the instrumental setup, the mobile phase used consisted of solvent A: Ultrapure water obtained from a Sartorius Arium Pro system containing 0.1 % formic acid (Cat. No. A117-50, Fisher Chemical) and solvent B: Acetonitrile (LC-MS Chromasolv, Cat. No. 15684740, Honeywell Riedel-de Haen) containing 0.1 % formic acid. The injection volume for all analyses was 50 µl and the flow rate was kept at 1 ml/min. The mobile phase gradient used was:

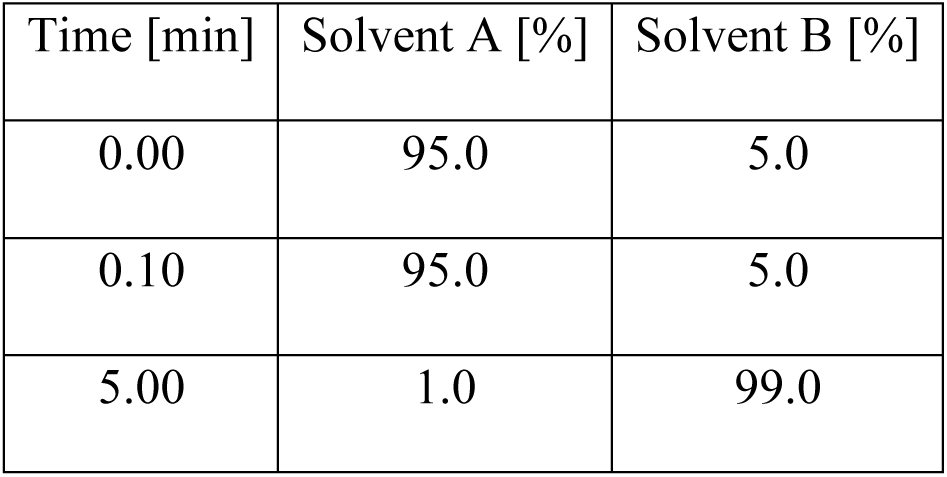
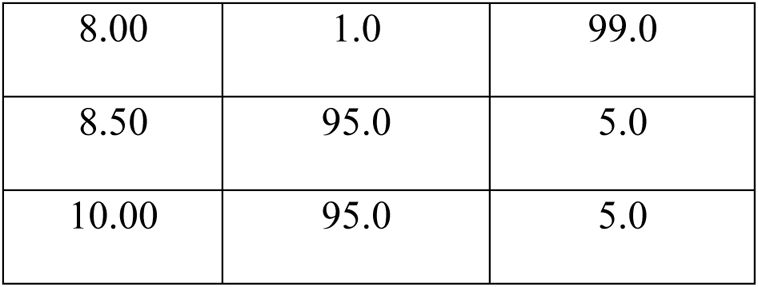

Using the results from the linear regression of the internal calibration measurement points (Supplementary Table 2) and the mean AUC values from the saturated aqueous solutions (Supplementary Table 3), the concentration C of the analytes in the saturated aqueous solution, expressed in µM, were calculated using equation (2):

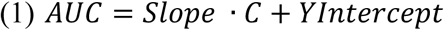

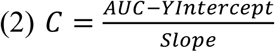

### Compound stability measurements in aqueous environment

The stability of the compounds was evaluated at a concentration of 12.5 µM in a 3:2 mixture of buffer (HEPES, 0.1 M, pH 7.4) and acetonitrile, with 1% DMSO. Acetonitrile was added because the concentration of 12.5 µM exceeds the thermodynamic solubility of the compounds in the buffer. This addition thereby prevented the formation of precipitates and a gradual decrease in compound concentration during the measurement. The resulting solution was stirred at 21 °C in a sealed vial (crimp neck vials, 5 mL, clear (Cat. No. 548-0611A) in combination with natural rubber septa (548-0565A) and aluminium crimp caps (548-3278A), VWR International) for 5 days. Samples of 100 µl were collected at 0, 24, 48, 72, 96, and 120 h and analysed using HPLC as described above. The AUC analysed was subsequently plotted as a function of the incubation time. (Supplementary Table 4)

### Computational studies

Computational studies of ligand binding to the AHR ligand binding domain (PAS-B) were performed using molecular docking. The experimental structure of the human AHR PAS-B domain (PDB ID: 7ZUB) ^20^, obtained with the cryo-electron microscopy (cryo-EM) technique, was taken from the Protein Data Bank. This structure includes the domain complexed with the indirubin ligand and other proteins that constitute the AHR cytosolic assembly. For docking calculations, non-AHR proteins were removed, and the resulting structure was pre-processed using Schrödinger Protein Preparation Wizard ^29^. Residue protonation states were assigned with PROPKA ^30^ at pH 7.0. Ligands of interest were prepared using the LigPrep module. Docking was performed with the Glide XP ^31^ (extra precision) algorithm included in the Schrödinger suite. The receptor grid for the AhR PAS-B domain was positioned in the centre of mass of the indirubin ligand derived from the cryo-EM structure ^20^.

### Graphing and statistical analysis

Experiments were performed with three or more replicates. Differences between two groups were analysed by Student’s t-test and multiple comparisons were determined by one-way ANOVA. If there were two factors (such as dose and time) investigated, data were analysed by two-way ANOVA followed by a post-hoc test. Data were expressed as means ± SEM, and P < 0.05 was considered significant. All analyses were performed using Microsoft Excel 2010 and GraphPad Prism 8.3.1 software.

## Results

### Imidazopyridines inhibit ER^+^ breast cancer cell proliferation via a mechanism independent of direct ERα binding

We previously identified six 2-phenyl-imidazo[1, 2α] pyridine derivatives that inhibited the proliferation of ER^+^ but not ER^-^ breast cancer cells. However, the mechanism of action of these compounds remains unclear ^17^. In this work, we first selected three of the six inhibitory compounds (X15695, X19724 and X19728) (Fig. 1A) and analysed their ability to inhibit the clonal expansion of MCF-7 breast cancer cells that express the wild-type or knock-in ERα mutations D538G and Y537S identified in endocrine-resistant breast cancer patients ^18^.

**Figure 1.**
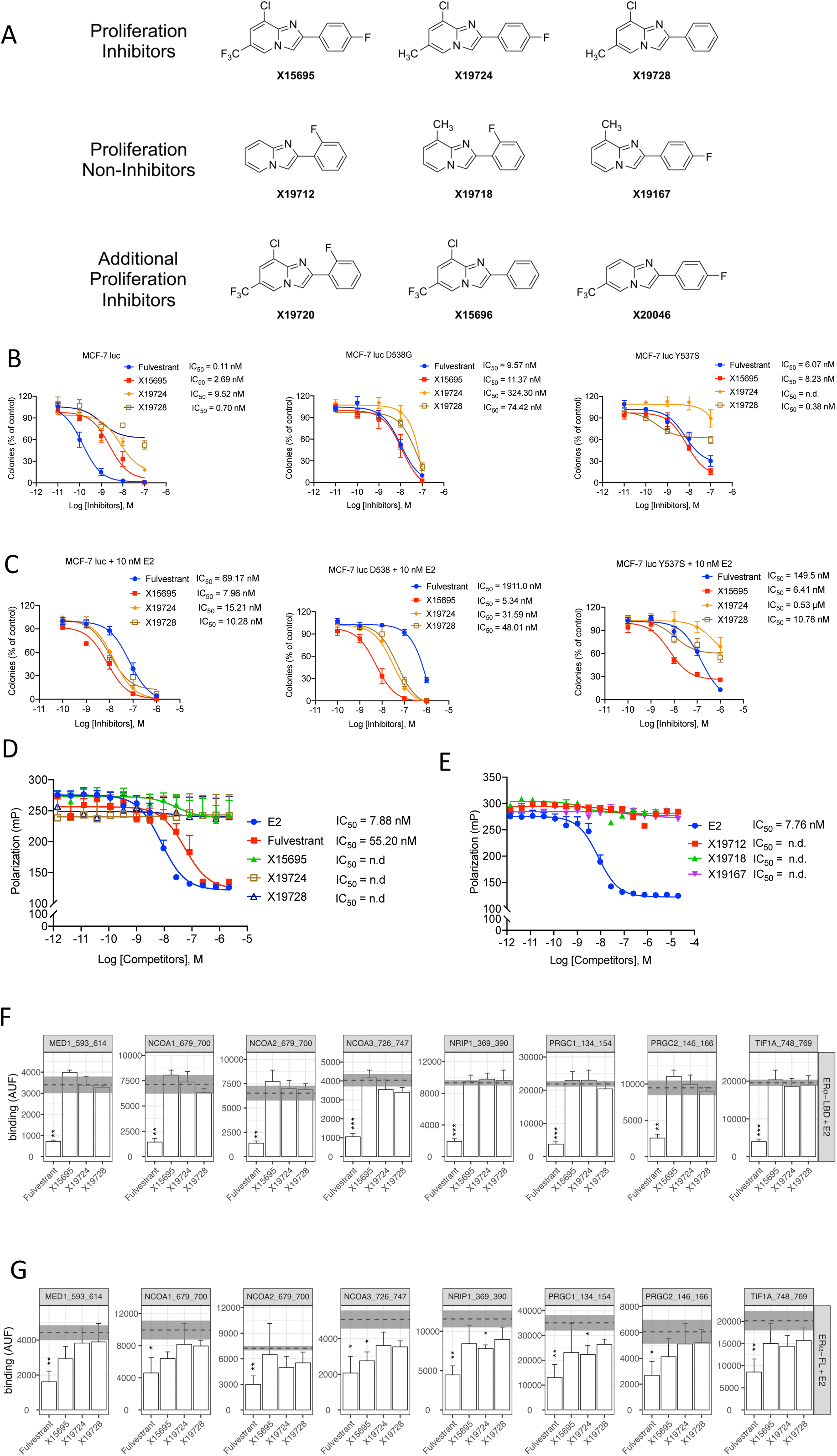
Effect of imidazopyridine derivatives on oestrogen receptor signalling. **A.** Structure of 2-phenyl-imidazo[1, 2a] pyridine derivatives used in this study and previously identified as inhibitors and non-inhibitors of ER+ breast cancer cell proliferation. **B and C**. Quantification of the concentration dependent action of the indicated compounds on the clonal expansion of the MCF-7 cells expressing the wildtype ER or the D538G of Y537S ER mutations (n = 3-6) in the absence **(B)** and presence **(C)** of 10 nM oestradiol. **D and E.** Competitive binding data generated using the PolarScreen™ ER Alpha Competitor Assay. Polarization values are plotted against the concentration of the test compound. Data were modelled using GraphPad Prism® software from GraphPad Software, Inc (n = 2 -8). **F and G**. Compound effect on E_2_-induced ER⍺ -coregulator interaction, zooming in on a subset of (well-established) ER-coregulators: MED1 (Mediator of RNA polymerase II transcription subunit 1); NCOA-1, -2, -3 (Nuclear receptor coactivator -1, -2, -3); NRIP1 (Nuclear receptor-interacting protein 1); PRGC1 (Peroxisome proliferator-activated receptor gamma coactivator 1a); PRGC 2 (Peroxisome proliferator-activated receptor gamma coactivator 1b) and TIF1A (Transcription intermediary factor 1-alpha). Recombinant ER⍺ LBD **(E)** or ER⍺ FL in MCF7 extracts **(F)** was treated with E_2_ EC_50_ (-8.5 logM) and subsequently incubated with 1000-fold excess (-5.5 logM) of each compound. Binding is represented as mean AUF (Arbitrary Units of Fluorescence of the detection antibody) +/-SEM (error bar in test compounds and grey area for control) of three technical replicates per indicated condition. Significance of test compound-induced modulation of ER control binding was assessed using Student’s t-Test (*p< 0.05; **< 0.01; ***< 0.001).

In colony forming assay with the three imidazopyridines, X15695 was identified as the optimal proliferation inhibitor in terms of potency (IC_50_ = 2.69 nM) and efficacy (Figs. 1B and 1C, left graphs). Although X19728 was somewhat more potent (IC_50_ = 0.70 nM), it was considerably less efficacious compared to X15695, as seen by its inability to effectively reduce colony number (Figs. 1B, graph on left). Note that potency refers to how much of a drug is needed to achieve a given effect, while efficacy is defined as the maximum effect a drug can produce, regardless of dose ^32^. In cells that express the wild-type ERα, the standard of care drug fulvestrant was more potent than X15695 (fulvestrant IC_50_ = 0.11 nM vs X15695 IC_50_ = 2.69 nM) but it is as potent as fulvestrant in cells that express the two mutant ERs (fulvestrant IC_50_ = 6.07 - 9.57 nM vs X15695 IC_50_ = 8.23 - 11.37 nM). Again, X19728 was potent (IC_50_ = 0.38 nM) but less efficacious in cells that express the mutant Y537S ERα (Fig. 1B, right graph). As the serum levels of premenopausal breast cancer patients undergoing anti-oestrogen therapy reach in some cases nanomolar levels of oestradiol (E_2_) (approx. 277.9 pg/ml or 1.02 nM) ^33^, we decided to evaluate the inhibition of clonal formation in the presence of E_2,_ to achieve a situation of fully liganded ERα. Intriguingly, the inhibitory action of fulvestrant compared to X15695 was significantly reduced in the presence of 10 nM oestradiol (E_2_) (Fig. 1C) varying in potency from 69.17 nM in cells expressing the wild-type receptor to the micromolar range (0.15 µM - 1.91 µM) in cells expressing the mutant receptors. On the other hand, X15695’s action remained relatively unchanged by E_2_ treatment (IC_50_ = 5 - 6 nM in the presence of E_2_ compared to 7 nM in the absence of E_2_) (Fig. 1C), indicating that the ERα may not be the primary target of X15695.

To confirm the observed differences in action of the imidazopyridine compounds and fulvestrant (a steroidal compound) in the presence of E_2_, we determined the relative binding affinities of these two classes of compounds to the ERα using a polar screen competition assay. This assay determines the effectiveness of ligands to compete with a selective fluorescent Fluormone tracer for binding to ERα. ERα and the Fluormone™ tracer form a complex, resulting in a high fluorescence polarization value. Compounds that bind ERα displace the Fluormone™ tracer from the complex and cause a decrease in fluorescence polarization. As expected, both oestradiol and fulvestrant efficiently displaced the Fluormone™ tracer from the ERα/Fluormone™ tracer complex. However, the three imidazopyridine derivatives did not (Fig. 1D), indicating that they do not associate with the ERα. Other compounds, X19712, X19718 and X19167 (Fig. 1A) from our collection of 2-phenyl-imidazo[1, 2α] pyridines that did not inhibit ER^+^ breast cancer cell proliferation ^17^ also failed to displace the Fluormone™(Fig. 1E).

We next determined whether the imidazopyridines that inhibited ER^+^ breast cancer proliferation (X15695, X19724 and X19728) functioned as ER antagonists using a nuclear receptor activity profiling (NAPing) assay. This assay measures nuclear receptor (NR) binding to a collection of coregulator-derived interactions motifs in vitro, and thereby mimics the recruitment of proteins which play a role in the canonical regulation of transcriptional activity. The oestradiol (E_2_)-activated ERα LBD or a full-length ERα was incubated with 101 peptides of well-known nuclear receptor coregulators immobilized on a solid support and the ability of the imidazopyridine compounds to disrupt the binding was determined at a single dose. Fulvestrant, used as a control in this assay, clearly disrupted binding of the coactivator peptides to either the liganded LBD or full length ER in a manner consistent with ER antagonism as previously reported ^34^ (Figs 1F and 1G; Supplementary Figs. 1A and 1B). In contrast, none of the imidazopyridine compounds showed this activity (Figs 1F and 1G; Supplementary Figs. 1A and 1B). These data support our hypothesis that the imidazopyridine derivatives do not directly target ERα. Nevertheless, in our previous RNA-seq and RT-PCR studies, X15695 attenuated ERα target gene expression in MCF-7 and T47D cells ^17^, indicating that it indeed has an effect on the receptor activity. We reassessed this action of X15695 and analysed the effect of the two other compounds (X19724 and X19728) on the transcription activity of ERα using a cell-based luciferase ER/ERE reporter assay. Attenuation of ERα transactivation was observed for all three compounds, albeit X19724 and X19728 actions were somewhat weaker (Fig. 2A). RT-PCR studies were also carried out with the three compounds on a select number of ERα target genes (PGR, TFF1, PDZK1 and GREB1) identified in our previous RNA-seq analysis ^17^. This assay also showed that all three compounds downregulated the expression of ERα target genes both in the absence and in the presence of E_2_ (Fig. 2B), although the action of X19728 was not significant in the presence of E_2_.

**Figure 2.**
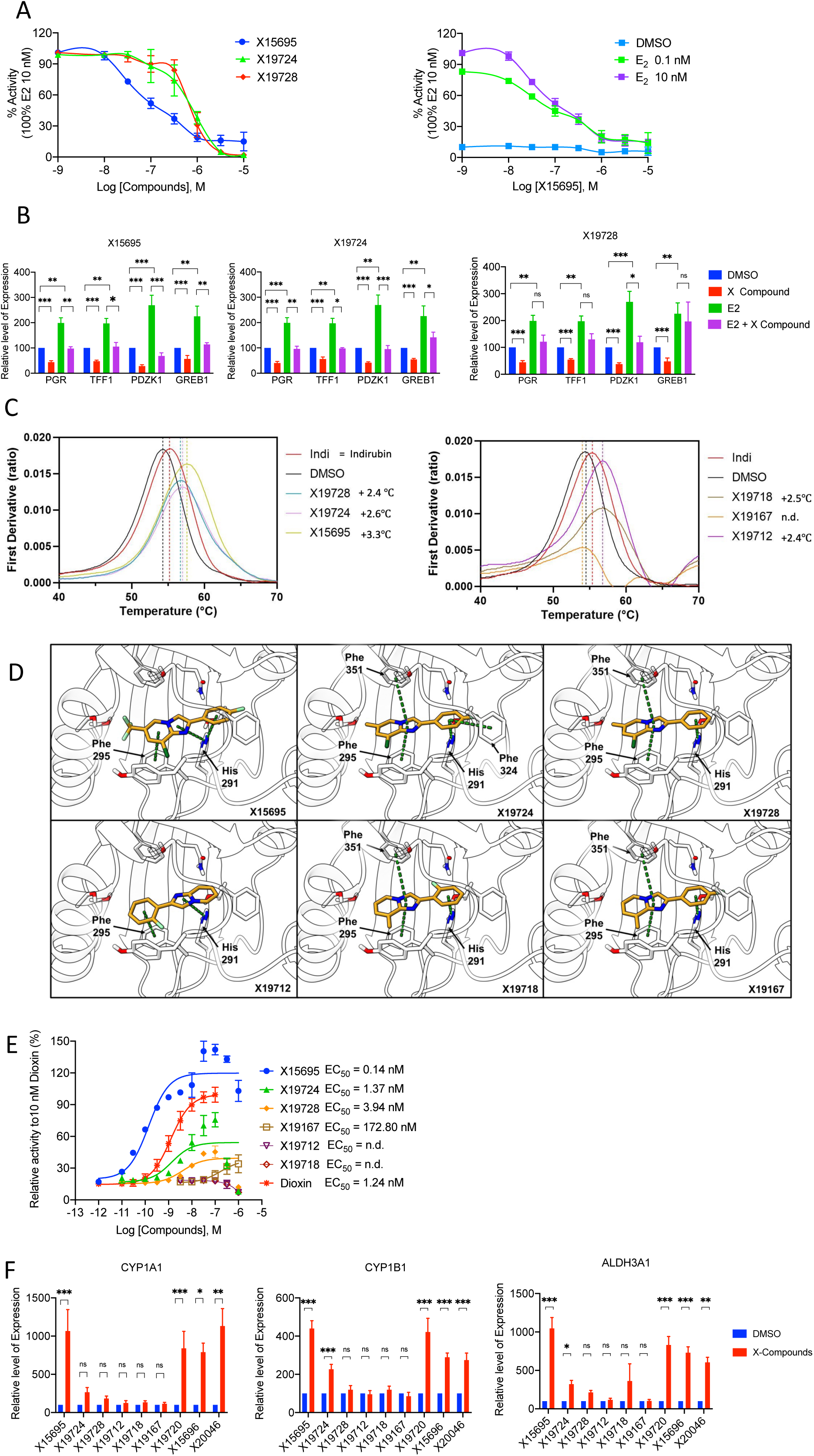
Binding of imidazopyridines to aryl hydrocarbon receptor and the resulting functional consequences. **A**. Results of reported gene assay on the action of the indicated imidazopyridines on the transactivation function of the ERα. Cells were treated with the indicated compounds (10 nM to 10 µM) in the presence of 10 nM (left part) or 0.1 or 10 nM (right part) E_2_ for 24 h. The results are the mean value ± SEM of independent biological replicates (n=2-4). **B.** Quantitative RT-PCR to detect the effect of the imidazopyridine derivatives on the expression of the indicated ER target genes in MCF-7 cells in the presence and absence of E_2_. Cells were treated with the indicated compounds (1 µM) in the presence and absence of 10 nM E_2_ for 24 h. The data represent the mean ± SEM (n = 3; (*p< 0.05; **< 0.01; ***< 0.001; ns is not significant). **C.** Nano-DSF analysis of the interaction between Hsp90-XAP2-AHR complexes and the indicated imidazopyridines. DMSO and Indirubin (indi) were used as negative and positive controls. The results are representative of independent biological replicates (n =3). **D**. Results of molecular docking calculations. The binding modes of the six compounds within the PAS-B binding domain of AHR are shown in a three-dimensional representation: ligands and the main interacting residues are shown as sticks, protein as white cartoons; π-π stacking interactions are highlighted with green dashed lines and the residues involved in these interactions are labelled. **E.** Results of reported gene assay on the action of the indicated imidazopyridines on the transactivation function of the AHR. The results are the mean value ± SEM of independent biological replicates (n=2-4). **F**. Quantitative RT-PCR to detect the effect of the imidazopyridine derivatives on the expression of the indicated AHR target genes in MCF-7 cells. Cells were serum starved for 72 h and treated with the indicated compounds (1 µM) for 16 h. The data represent the mean ± SEM (n = 3-12; *p< 0.05; **< 0.01; ***< 0.001; ns is not significant).

We hypothesised that since the imidazopyridines do not bind ERα, they most likely attenuate ERα gene expression indirectly, possibly through association with a component that in turn binds ERα. To identify such a component, we re-analysed our previous RNA-seq experiments carried out in MCF-7 and T47D cells after treatment with E_2_ and X15695, where xenobiotic metabolism was identified as one of the topmost signalling pathways in the Gene Set Enrichment Analysis (GSEA) ^17^. Heatmaps of the Log_2_ fold-change in gene expression in X15695 vs vehicle or E_2_ + X15695 vs E_2_ treatment in MCF-7 and T47D cells identified several genes involved in xenobiotic metabolism (Supplementary Figs. 2A and 2B). Importantly, the transcriptional activity of these genes is controlled by the aryl hydrocarbon receptor (AHR) which is known to interact with and alter the transcriptional activity of, among others, the oestrogen and androgen receptors ^35^. We therefore postulated that our imidazopyridine derivatives possibly function by interacting with the AHR.

### The Aryl Hydrocarbon Receptor is the direct molecular target of antiproliferative imidazopyridines

Using differential scanning fluorimetry (nano-DSF) that monitors the thermal stabilisation of proteins upon ligand binding, we confirmed that X15695, X19724 and X19728 bound purified AHR-Hsp90-XAP2. Notably, they stabilised the AHR to a greater degree than the classical AHR ligand, indirubin (Indi), which was used as a positive control in these assays (Fig. 2C). The ranking order was X15695 (+3.3°C) > X19724 (+2.6°C) > X19728 (+2.4°C) (Fig. 2C, left panel). Other 2-phenyl-imidazo[1, 2α] pyridine derivatives X19712 and X19718 (Fig. 1A) that poorly inhibited breast cancer cell proliferation ^17^ also induced thermal shifts (X19712: +2.4°C, X19718: +2.5°C) (Fig. 2C, right panel) while another compound X19167 (Fig. 1A) that also poorly inhibited breast and prostate cancer cell proliferation ^17^ showed no stabilisation of the AHR and it likely does not interact with the AHR (Fig. 2C, right panel). These findings suggest that the inhibition of cell proliferation by the imidazopyridines may not always be correlated with interaction with the AHR.

We next sought to gain more detailed insight into the interactions, including binding modes and potential binding affinities of the compounds employed in the nano-DSF studies, using *in silico* docking calculations of the compounds in the AHR ligand binding domain. For this, we used the relevant human experimental structures of the AHR PAS-B domain (PDB IDs 7ZUB) in complex with indirubin ^20^. We utilized the Glide software, a part of the Schrödinger suite, for all docking calculations. As represented in Fig. 2D, most compounds formed a π-π stacking interaction network (represented by green dashed lines). This network was primarily anchored by a stacking interaction between the imidazopyridine ring of the ligands and the aromatic side chain of Phe295, with a T-stacking further stabilising the secondary phenylic ring. Additional ππ stacking interactions were observed with Phe351 and Phe324. The docking scores showed that X15695 has the highest predicted affinity (the most negative score) while X19724 and X19728 exhibited slightly lower affinities (Table 1). The remaining three ligands, that did not inhibit ER^+^ breast cancer proliferation had the lowest predicted binding affinities (less negative docking scores).

**Table 1.** Glide score and predicted PKa of different imidazopyridine compounds.

| Compounds | Inhibition of Proliferation | Glide XP Score (kcal mol<sup>-1</sup>) | Predicted pKa |
| --- | --- | --- | --- |
| X15695 | Yes | -10.13 | 3.09 |
| X19724 | Yes | -9.357 | 4.50 |
| X19728 | Yes | -8.791 | 4.69 |
| X15696 | Yes | -10.055 | 3.28 |
| X19720 | Yes | -10.377 | 3.28 |
| X20046 | Yes | -9.76 | 4.83 |
| X19712 | No | -8.354 | 6.27 |
| X19718 | No | -8.608 | 6.55 |
| X19167 | No | -8.65 | 6.36 |
| imidazo(1,2-a)pyridine |  |  | 6.69 |

To complete the AHR binding analysis for all the six imidazopyridines previously identified as potent cell proliferation inhibitors ^17^ (Fig. 1A), docking calculations were also performed on the remaining three compounds in that group (X19720, X15696, and X20046). These three compounds exhibited docking scores comparable to that of X15695 (Table 1), indicating high binding affinities for the receptor. Also, their binding modes, predicted by docking calculations (Supplementary Fig. 3), were like those previously observed for the other inhibitors studied in this work (Fig. 2D).

Intriguingly, during the preparation phase of the compounds, we noted that the aqueous phase pKa values (predicted by the *Epik* tool) present important differences among the compounds (Table 1). All the six experimentally active compounds that inhibited breast cancer cell proliferation (X15695, X19724, X19728, X15696, X19720, X20046) were predicted to be almost exclusively neutral at physiological pH, with pKa values for the imidazopyridine nitrogen ranging from 3.5 to 4.8. In contrast, all experimentally inactive compounds (X19712, X19718, X19167) had a pKa close to 6.5, implying they exist as an equilibrium mixture of neutral (∼89%) and protonated (∼11%) forms. As a control, we also predicted the pKa for the non-functionalized imidazo[1,2-α]pyridine compound, obtaining a value of 6.69, which aligns well with the experimental value of 6.79 reported in the IUPAC Digitized pKa Dataset ^36^, thus confirming the reliability of our approach. Two effects may contribute to the observed behaviour of differently substituted imidazopyridine rings. On the one hand, the electron-withdrawing substituents that lead to a low pKa also render the aromatic system electron-deficient, enhancing the crucial π-π stacking interactions with key residues like Phe295 and His291, as supported by our docking calculations. On the other hand, a high pKa is detrimental to binding affinity. This might seem counterintuitive, as the resulting protonated cation would be expected to form even stronger cation-π interactions. However, the predominantly hydrophobic nature of the pocket imposes a large energetic penalty for the desolvation of a charged species, making the binding of the protonated form highly unfavourable from a thermodynamic point of view. Therefore, we propose that potent AHR ligands must first exist in a neutral state to efficiently partition into the binding site. This condition is met only by the compounds with low pKa values, which combine favourable neutrality with an electronic profile optimised for π-π stacking.

### Imidazopyridines act as potent agonists of the AHR signalling pathway

To determine whether ligand interaction with the AHR correlates with functional activity of this receptor, we first analysed the transactivation function of the AHR in a cell-based luciferase reporter gene assay after treatment of the cells with the first two groups of compounds (Fig. 1A). In this assay, X15695 upregulated the transcriptional activity of the AHR with potency that surpassed that of the classical AHR ligand dioxin, whereas X19724 and X19728 showed only modestly enhancement (Fig. 2E). Their effect was marginal both in terms of potency and efficacy. The other imidazopyridine compounds that showed reduced binding to the AHR were functionally inactive in the reporter gene assay at the concentrations tested (Fig. 2E).

The AHR transactivation assay was further extended to cover all the compounds in this study (Fig.1A) employing RT-PCR gene expression analysis of AHR target genes CYP1A1, CYP1B1 and ALDH3A1 both in MCF-7 (Fig. 2F) and T47D cells (Supplementary Fig. 4). Consistent with reports on the behaviour of AHR ligands ^37^, a gene and cell line-selective effect was evident with the action of the compounds X19724 and X19728 (Fig. 2F and Supplementary Fig. 4). In contrast, X15695, X15696, X19720 and X20046 were more consistent in their effects and enhanced the transcriptional activity of the AHR target genes in both MCF-7 and T47D cells (Fig. 2F and Supplementary Fig. 4). Compounds X19712, X19718 and X19167 that poorly inhibited breast cancer cell proliferation failed to activate AHR target gene expression in the two cell lines (Fig. 2F and Supplementary Fig. 4). Further characterisation of two representative members of the subgroup that enhanced the transcriptional activity in both cell lines (X19720 and X15696) showed that, like X15695, they outperformed fulvestrant in the inhibition of proliferation of MCF-7 cells in the presence but not in the absence of E_2_ (Supplementary Fig. 5). Other descriptors of drug design used to further characterise three members (X15696, X15695 and X19720) of this subgroup showed that they have marginal thermodynamic solubilities reaching 5 µM, 3 µM and 1.4 µM respectively using the shake flask method (materials and methods and Supplementary Tables 2 and 3). They were nonetheless found to be remarkably stable over 5 days at 21 °C with no obvious decomposition detected (Supplementary Table 4). These findings need to be taken into consideration in future development and formulations of these compounds for therapeutic applications.

### The antiproliferative effect of imidazopyridines is strictly dependent on the AHR

To determine to what degree the affinity of the compounds for the AHR is correlated with the inhibition of proliferation of ER^+^ breast cancer cell lines, we blocked the activity of AHR with CH223191 ^38^ and then tested whether the compounds could still inhibit colony growth in MCF-7 cells. Except for X19728 that had no effect on colony formation (Supplementary Fig. 6, red bars), the inhibition of colony expansion by the compounds X15695, X15696, X19720 and X20046 (and, to a lesser extent, X19724) was clearly abrogated by the AHR antagonist CH223191 (Supplementary Fig. 6, compare red with purple bars). Fulvestrant clearly inhibited colony growth in the presence of CH223191, demonstrating its independence from AHR and a different mode of action for the inhibition of ERα-dependent breast cancer cell growth.

As the competitive inhibitor CH223191 leaves the AHR intact and only blocks its activity, we also generated a complete AHR knock-out (KO) in MCF-7 and T47D cells using the clustered regularly interspaced short palindromic repeats (CRISPR) technology that permanently removes AHR from cells (Supplementary Fig. 7A). Two MCF-7 (#2 and #17) and T47D (#7 and #30) AHR KO clones (Supplementary Figs. 7B and 7C) showed decreased proliferation compared to the empty vector transfected cells, demonstrating that the AHR contributes to the proliferation of these cells (Fig. 3A and Supplementary Fig. 8A). When treated with the imidazopyridines, the MCF-7 AHR KO clones became resistant to the inhibition of colony growth that is normally observed in the control vector transfected cells (Fig. 3B). These findings agree with the results obtained following AHR inhibition with CH223191 (Supplementary Fig. 6) showing an even stronger effect upon CRISPR-mediated ablation of AHR (Fig. 3C). The inhibitory action of X19728 in MCF-7 cells was again less pronounced as opposed to X15695 (Fig. 3C). Similar results were obtained in T47D KO clones (Supplementary Figure 8B and 8C), but, overall, the T47D cells were less sensitive than MCF-7 cells to inhibition by the X compounds (Supplementary Figure 7B). In both cell types, X15695 was the most effective compound in reducing cell proliferation.

**Figure 3.**
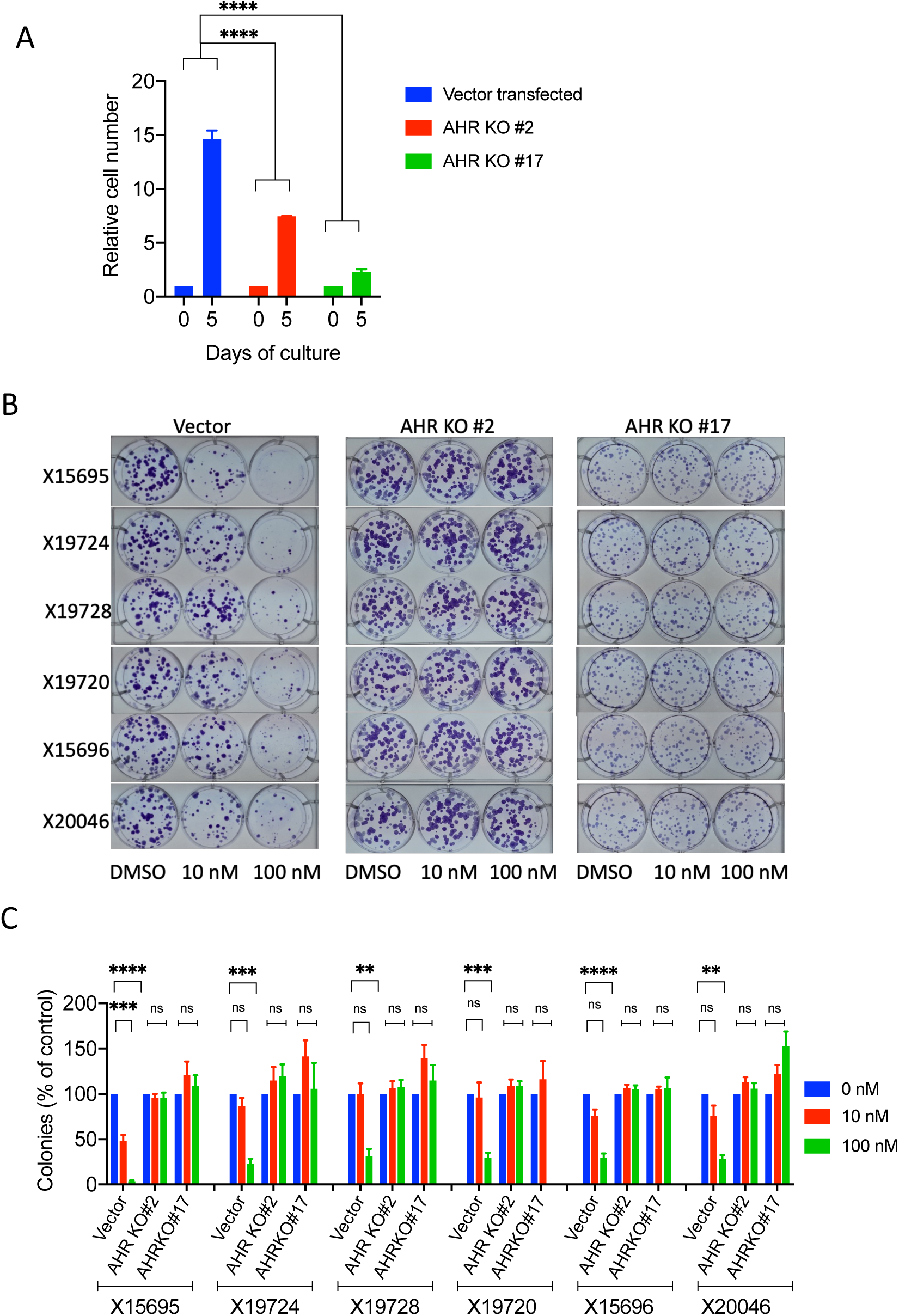
2-phenyl-imidazo[1, 2a] pyridine-mediated inhibition of ER+ MCF-7 breast cancer cell proliferation via AHR. **A**. Proliferation of empty vector transfected MCF-7 cells and MCF-7 AHR KO clones #2 and #17. Cells were cultured in complete media and counted on days 0 and 5. The starting number of cells was 1×10^4^ per well. Data are the averages of three independent experiments ± SEM, normalized to day 0., ****p ≤ 0.0001. **B**. Representative images of the clonal expansion of empty vector transfected MCF-7 cells or AHR CRISPR-CAS9 KO MCF-7 cell clones #2 and #17 after treatment with the indicated concentrations of X compounds. **C.** Quantification of the concentration dependent action of the indicated compounds on the clonal expansion of empty vector transfected MCF-7 cells and AHR KO MCF-7 clones #2 and #17. The results are shown as bar charts and they represent the means ± SEM, n = 3-4. *** p < 0.001; **** < 0.0001; ns is not significant.

### AHR activation by imidazopyridines triggers the proteasomal degradation of ERα

We have previously shown that the inhibition of ER^+^ cell proliferation by X15695 is associated with a decrease in ERα stability. To determine whether destabilisation of ERα is unique to X15695, we investigated whether the other X compounds that reduced ER^+^ cell proliferation also altered ERα stability. We treated MCF-7 cells for 48 h with increasing concentrations of all six imidazopyridines that we previously identified as good inhibitors (namely X15696, X19720, X15695, X20046, X19728, X19724), in addition to the compounds X19712, X19718 and X19167 that we identified as poor inhibitors of breast cancer cell proliferation (Fig. 1A) ^17^. Analyses of their ability to degrade ERα using an immunoblot assay and quantification of the results identified five compounds (X15695, X19724, X19720, X15696 and X20046) as being the most proficient in destabilising ERα (Figs. 4A and 4C). X19728, like the compounds X19712, X19718 and X19167 that poorly inhibited breast cancer cell proliferation ^17^, showed no significant ERα destabilisation at the concentrations used in the study (Figs. 4B and 4C). This was not surprising considering X19728 showed the weakest effect in our studies.

**Figure 4.**
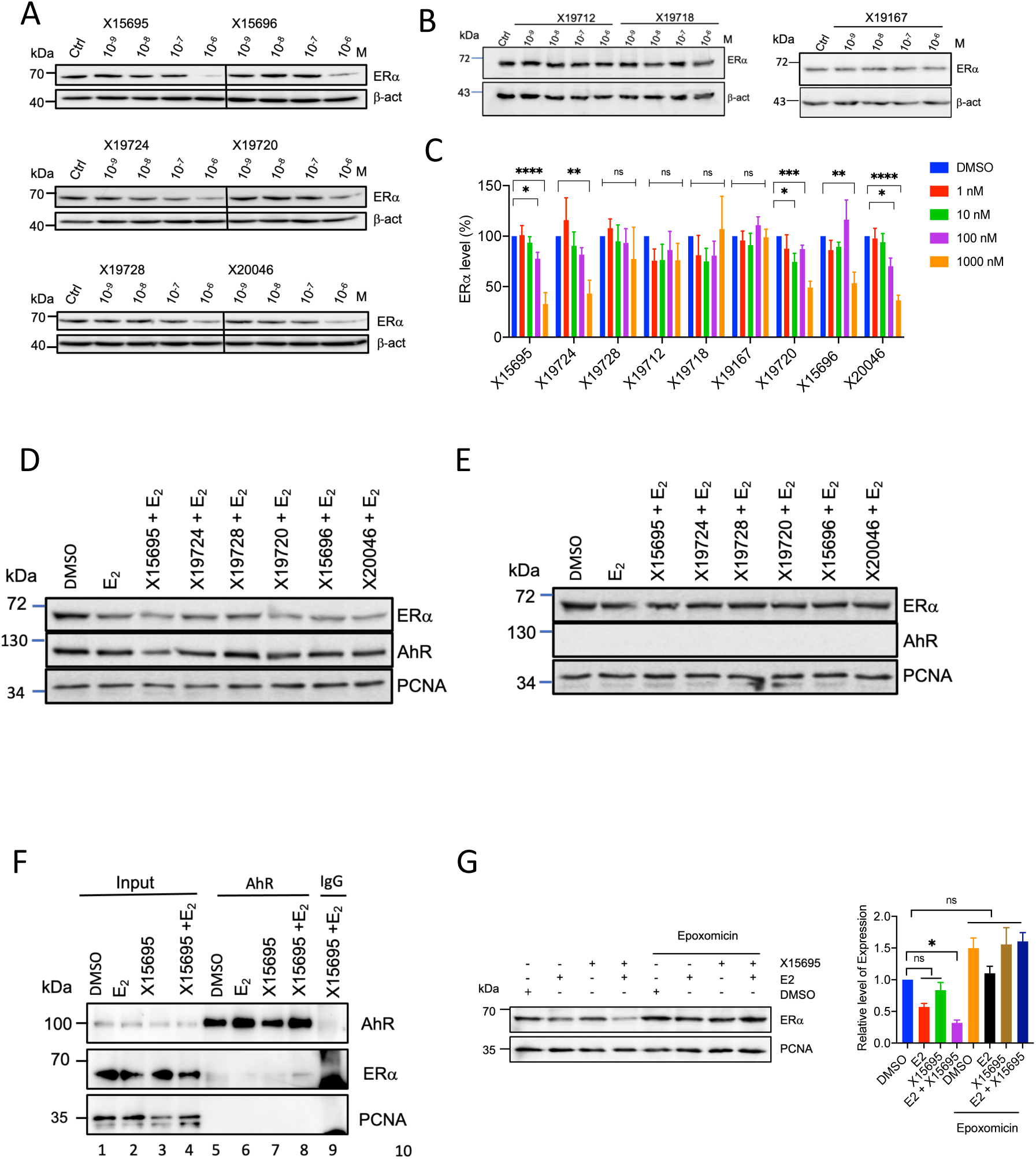
Degradation of ERa by X15695 is mediated via the proteasome A. and. **B.** Western blot analysis of ER7 after treatment of MCF-7 cells with the indicated concentrations and imidazopyridine compounds for 48 h with anti - ER7 antibody and an anti-β-actin antibody that was used for the loading control. **C.** Protein signals in the Western blots in **(A**) and **(B**) were quantified. The bar charts show the level of ERa expression relative to the β-actin expression in **(A)** and **(B)** and they represent the means ± SEM (n = 3 - 6. *p< 0.05; **< 0.01; ***< 0.001; ****< 0.0001; ns is not significant)**. D**. MCF-7 cells (empty vector transfected) and **E.** MCF-7 AHR KO clone #2 were cultured in hormone deprived medium for 3 days and treated for 8 h with E_2_ (10 nM) or E_2_ + the indicated X compounds (1 µM). Western blots were carried out with lysates of these cells using anti-ERa and AHR antibodies along with anti-PCNA antibody to demonstrate equal protein loading. **F.** Western blot analysis of ERa using immunoprecipitated extracts of MCF-7 cells treated with DMSO, E_2_ (10 nM), X15695 (1 µM) and E_2_ +X15695 for 1h. 70 µg of cell extracts was used as input control while 1,900 µg was immunoprecipitated with anti-AHR or IgG antibodies immobilized on protein A/G agarose beads. **G.** Western blot analysis of ERa in extracts of MCF-7 cells pre-treated with 4.5 μM Epoxomycin for 1 h in the presence and absence of DMSO, then E_2_ (10 nM), X15695 (1 µM) and E_2_ +X15695 were added and incubated for further 2.5 h. Anti-PCNA antibody was used for the loading control. The bar chart (right) shows the results of the level of expression of ERa protein relative to PCNA and they represent the mean ± SEM (n=3; *p< 0.05; ns refers to non-significant result).

To determine to what extent the AHR plays a role in the degradation of the ERα, we compared the action of the six ligands in the empty vector transfected MCF-7 and the MCF-7 AHR KO #2 cells after treatment for 8 h. While these compounds destabilised ERα to varying degrees consistent with their affinity for the AHR (Fig. 4D), they did not have any effect on the level of the ERα in the AHR KO cells (Fig. 4E), demonstrating that the X compound-induced destabilisation of ERα is dependent on its interaction with AHR.

We next sought to find out how the degradation of the ERα occurred and investigated whether the AHR and the ERα interact directly. In co-immunoprecipitation studies, we showed that ERα and AHR formed a complex as early as 1 h after treatment with E_2_ and X15695 (used as a prototype of the inhibiting compounds) (Fig. 4F, lane 8). Furthermore, in the presence of the selective proteasome inhibitor epoxomicin ^39^, X15695-mediated destabilisation of ERα was abrogated, indicating that the ERα degradation occurs via the proteasome (Fig. 4G). These results identify X15695 as a member of a distinct family of imidazopyridines that targets the AHR to inhibit ER^+^ breast cancer cell proliferation.

### X15695 inhibits the growth of organoids from patient-derived xenograft tumours (PDxO)

Our previous preclinical studies showed that X15695 inhibited the growth of MCF-7 tumour xenograft in mice ^17^. To further assess its clinical potential, we employed patient-derived organoids (PDxOs), generated from ER^+^ breast tumours previously established as patient-derived xenografts ^19^. PDxOs preserve critical features of the original tumours, including genetic and epigenetic profiles, as well as aspects of tissue architecture and cellular heterogeneity. This makes them robust and clinically relevant models for predicting therapeutic responses ^40^. We used previously characterised PDxO ^19^ established from a PDX of a treatment naïve ER^+^/HER2^-^ breast cancer patient. PDxOs were embedded in 3D Matrigel for 24 h and then treated with increasing doses of fulvestrant or X15695 for 7 days. Viability assay showed that X15695 significantly reduced PDxO survival in a dose-dependent manner, whereas the response to fulvestrant was more modest (Fig. 5A and B). These results support the efficacy of X15695 in targeting ER⁺ breast cancer and underscore its potential as a therapeutic agent for further preclinical and clinical development.

**Figure 5.**
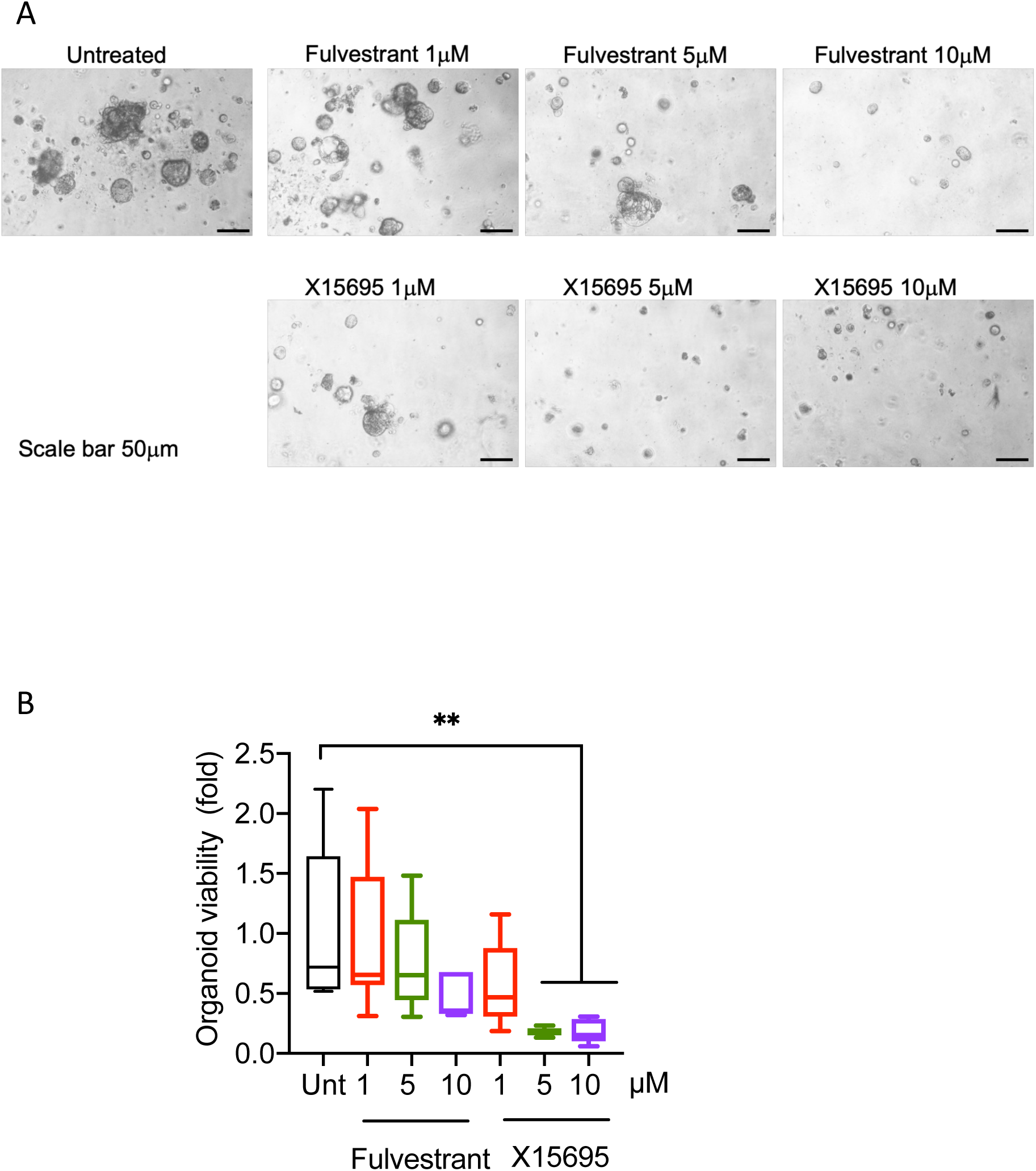
X15695 reduced the viability of ER^+^ breast cancer patient-derived models (PDxO) **A.** Representative contrast phase images of PDxOs untreated or treated with fulvestrant or X15695 at the indicated doses, corresponding to conditions quantified in panel **(B)**. Images were acquired using a 4X objective. Scale bar is 50mm. **B.** Graph reports the viability of PDxOs treated with increasing concentrations of fulvestrant or X15695 for 7 days, measured using the CellTiter-Glo assay. PDxOs were cultured in 3D Matrigel; drug-containing media were refreshed on day 4. Data are presented as fold change relative to the untreated control). Statistical significance was assessed by one-way ANOVA test (p < 0.05). Box plots show median, minimum, and maximum values; individual data points represent independent experimental replicates. A minimum of six replicates was used per condition (N=6-8). **< 0.01.

## Discussion

The ERα is a critical target of therapeutic significance in the control of ER^+^ breast cancer. However, the clinically available ER antagonists used for the treatment of breast cancer have limitations due, in part, to mutation of ERα gene (*ESR1*) that make the tumours intractable to treatment. SERDs have therefore been developed to overcome these problems but fulvestrant, the first clinically available SERD, showed only modest inhibition of mutant ERα, compared to wildtype ERα ^3^. Although other SERDs have since been developed or are in development, it remains important to develop further SERDs especially those with a different mode of action for the effective inhibition of breast cancer cells with ERα mutations.

We have previously identified the compound X15695, a derivative of 2-phenyl-imidazo[1, 2α] pyridine, as a SERD but its mode of action has not been further investigated. Here, we have characterized the action of X15695 and additional derivatives in this compound class. Compared to the standard of care compound fulvestrant, the imidazopyridine derivatives in our study do not interact with ERα nor do they inhibit NR-box (LxxLL)-mediated recruitment of coactivators that is otherwise required for ERα agonism or targeted by most conventional SERDs or SERMs. Instead, the imidazopyridines reported here, interact with the AHR, which leads to degradation of ERα. Pharmacological inhibition of the activity of AHR or genetic ablation of the AHR attenuates the inhibitory action of the compounds on breast cancer cell proliferation, demonstrating that these anti-cancer effects are dependent on functional AHR in cells.

In this work, we have identified a subgroup of four imidazopyridine compounds including X15695, which inhibit the proliferation of MCF-7 cells as potently as fulvestrant. Moreover, these compounds fully retain their inhibitory function in cells that express the two main clinically relevant ERα mutants D578G and Y5237S. Importantly, in the presence of oestradiol, a condition that could mirror the pre-menopausal state of some ER^+^ breast cancer patients, these compounds outperform fulvestrant in inhibiting the proliferation of MCF-7 cells expressing either the wild-type or mutant ERα, and PDxOs from a ER^+^ breast cancer patient that are grown in 3D in oestradiol-containing medium. We further showed that these imidazopyridines interact with the AHR and they need the AHR for their inhibitory action of ER^+^ breast cancer cells.

The AHR belongs to the basic helix-loop-helix-PER-ARNT-SIM (bHLH-PAS) family of transcription factors, characterised by a conserved bHLH DNA binding domain at its N-terminus, followed by tandem PAS domains (PAS-A and PAS-B), and a transactivation domain ^41^. The PAS-B domain is the primary site for binding of small-molecule ligands to the AHR ranging from polycyclic aromatic hydrocarbons or polychlorinated biphenyls but the precise basis for these diverse ligand interactions has so far remained elusive ^37^. Recently, the crystal structures of the AHR in complex with hydrocarbon receptor nuclear translocator (ARNT) and DNA bound to each of six established AHR ligands have been solved. This revealed that for transcriptional activity, the PAS-B domain utilizes eight conserved residues whose dynamic rearrangements account for the ability to bind to ligands through hydrophobic and π-π interactions ^37^.

Apart from the regulation of transcription, liganded AHR has non-genomic action such as binding to and degrading transcription factors such as the ERα and the androgen receptor (AR) ^42–46^. However, the features of the ligands that determine these effects have also remained elusive and the crosstalk of the AHR with the ERα is controversial. While some AHR ligands are reported to induce ERα degradation other ligands negatively regulate activity independent of degradation whereas yet other ligands are reported to activate ERα signalling ^47,48^. Other studies report that the AHR-mediated regulation of ERα transactivation occurs in a gene selective and cell-context specific manner ^48^. Ligands including 2,3,7,8-tetrachlorodibenzo-*p*-dioxin, 6-formylindolo(2,3 b) carbazole, carbidopa and the prenylflavone icaritin ^42–46,49^ have been described to endow the AHR with the ability to degrade ERα but no structure-activity relationship studies have been carried out.

In the present study we have used 2-phenyl-imidazo[1, 2α] pyridine derivatives with different affinities to the AHR in binding and computational docking experiments to determine which ligands inhibit ER^+^ breast cancer cell proliferation and destabilise the ER and which do not. We noted that 2-phenyl-imidazo[1, 2α] pyridine derivatives that strongly interact with the AHR most potently inhibit ER^+^ breast cancer cell proliferation and degrade the ER. These compounds were predicted to have low pKa values, suggesting the presence of electron-withdrawing substituents that deplete electron density from the aromatic ring. Decreasing the electron density in the π-cloud causes a reduction of the electrostatic repulsion between π-systems, thereby enhancing the π-π stacking interactions as has been described in several theoretical models ^50–52^. It is thus conceivable that ligands bearing electron-withdrawing substituents exhibit enhanced AHR affinity due to an extra-stabilization of the π-stacking interactions with the aromatic residues in the binding cavity, particularly with F295. These properties are clearly identifiable in our four compounds X15695, X15696, X19720 and X20046.

The imidazopyridines analysed in this work have also been reported to inhibit the proliferation of AR^+^ prostate cancer cells in our previous study ^17^. In those studies, the inhibition of prostate cancer cell proliferation was linked to the destabilisation of the wild-type AR in LNCaP cells or the AR wild-type and AR V7 splice variant in 22Rv1 prostate cancer cells ^17^. Although the regulation of AR action is not the topic of this study, it is important to mention that destabilisation of the AR by liganded AHR has previously been described ^53–55^ and it is likely that a mechanism similar to what we have described here for ER^+^ breast cancer cells may also apply for the inhibition of AR^+^ prostate cancer cell proliferation.

The exact mechanism of how the AHR destabilises nuclear receptors or other proteins is not clear. One possibility is that the AHR acts as a substrate-recognition subunit to recruit ERα/AR for proteolysis by assembling a ubiquitin ligase complex, CUL4B(AHR) ^42,56,57^. The involvement of another E3 ligase has also been reported. For example, indole-3-carbinol-dependent activation of AHR has been shown to initiate the E3 ubiquitin ligase ring box 1 (Rbx-1) and proteasomal degradation of ERα ^58^. Evidence that E3 ligase complexes recruited by the AHR bring about protein degradation comes from recent studies that show that the incorporation of AHR ligands into proteolysis-targeting chimaeras (PROTACs) generate molecules for the targeted degradation of cellular proteins ^59^. Other proposed putative actions of AHR for the inhibition of ERα signalling pathway include the attenuation of ER target gene expression by the direct binding of the activated AHR and ARNT heterodimer complex to inhibitory xenobiotic response elements (iXRE) in ER target genes. Alternatively, squelching of shared coactivators, including ARNT and synthesis of an unknown inhibitory protein have been proposed ^60^.

Although our results show that the imidazopyridines target AHR for their anti-proliferative action, our previous studies have also shown the involvement of p53 signalling in the action of the imidazopyridines. X15695 was shown to activate p53 and knock-down of p53 partly attenuated the inhibitory action of X15695 on MCF-7 and T47D cell proliferation ^17^. It is therefore likely that AHR and p53 cooperate to inhibit proliferation of the breast cancer cells upon treatment with the imidazopyridine derivatives. Future studies will be needed to decipher the intricacies of these signalling pathways. At the moment, our findings identify the AHR as a key component in the action of the 2-phenyl-imidazo[1, 2α] pyridine derivatives and demonstrate the features on these ligands needed for the AHR-mediated inhibition of proliferation of ER^+^ breast cancer cells.

## Acknowledgements

We thank Carsten Weiss for fruitful discussions and suggestions. We acknowledged with thanks the help we received from Celine Moser and Claudia Muhle-Goll with the use of the Tecan Reader for the fluorescence polarisation studies. M.P. received funding for his contribution to this work from the China Scholarship Council (CSC), grant No. 201807090113; B.B. received funding from Ministry of Health (RF-2021-12371961) and AIRC (IG20061). This project was supported by the core facility “Molecule Archive” of the German Research Foundation (Deutsche Forschungsgemeinschaft, DFG project number: 284178167).

## Authors’ Contributions

A.C.B.C. conceived and directed the study. C.B. performed RT-PCR experiments and clonogenic assays together with K.K. and M.R., Z.W. and M.R. performed the cell proliferation studies. C.B., J.M. and J.S. performed Western blot assays. S. M. and L.B. performed docking and computational analysis. M.P. generated material and analysed RNA-seq datasets. R.H. carried out NAPing experiments. C.W.G. performed solubility and stability experiments of the compounds. L-W.L. and S.K.K. characterised some of the compounds in reporter gene assay and performed clonogenic experiments in AHR KO cells. S. S. and W. B. performed the nano-DSF experiments. M.G. and P. B. carried out reporter gene assays. L.B. and S.A. provided MCF-7 mutant cells and information on how to use them. K.K. and M.R. performed the polar screen assay and analyses of fluorescence polarisation. S.B., N.J and S.B. provided the compounds for the study. I.S. and B.B generated organoids from PDX tumours and analysed the action of the X compounds on them. G.D. was involved in conceptualization, provision of resources, formal analyses and supervision. A.C.B.C., S.M. and L.B. wrote the paper with critical feedback from all the coauthors.

## Supplementary Figure legends

**Supplementary Figure 1**. Effect of X compounds on E_2_-induced ER⍺-coregulator peptide interaction. **A.** Recombinant ER⍺ LBD or **B.** ER⍺ FL in MCF7 extracts was partially treated with E_2_ EC_50_ (-8.5 logM) and subsequently incubated with 1000-fold excess (-5.5 logM) of each compound for 30 min. Each panel shows plots of the level of ER binding in the absence of compound (ctrl, E_2_ only, black line) and presence of the test compounds (coloured line, see legend) to each of 101 coregulator-derived NR-binding motifs on the NAPing platform. Binding is represented as mean of three technical replicates per indicated condition. Significance of test compound-induced modulation of ER control binding was assessed using Student’s t-Test (*p<.05;**<0.01;***<.0001).

**Supplementary Figure S2**

Hallmark gene analysis showing genes in the xenobiotic metabolism signalling pathways targeted by X15695. **A**. Gene Set Enrichment Analysis (GSEA) plots of xenobiotic metabolism signalling pathway identified as one of the topmost gene sets in a comparison of X15695 vs vehicle and E_2_ + X15695 vs E_2_ in MCF-7 and T47D datasets. **B.** Heatmaps of Log_2_ fold-change in gene expression in the comparison of X15695 vs vehicle and E_2_ + X15695 vs E_2_ treatment of MCF-7 and T47D cells.

**Supplementary Figure S3**

Results of molecular docking calculations. The binding modes of the three compounds within the PAS-B binding domain of AHR are shown in a three-dimensional representation: ligands and the main interacting residues are shown as sticks, protein as white cartoons; π-π stacking interactions are highlighted with green dashed lines and the residues involved in these interactions are labelled.

**Supplementary Figure 4**

Effect of X compounds on AHR target gene expression in T47D cells. Quantitative RT-PCR carried out to detect the effect of the indicated imidazopyridine derivatives on the expression of three AHR target genes in T47D cells. Cells were hormone-starved for 72 h and treated with 1 µM of the indicated compounds for 16 h. The data represent the mean ± SEM (n = 3-12; *p< 0.05; **< 0.01; ***< 0.001; ns is not significant.).

**Supplementary Figure 5**

Comparison of the action of imidazopyridine derivatives and fulvestrant in clonal expansion of MCF-7 cells expressing wild-type and mutant oestrogen receptors. **A, B**. Quantification of the concentration dependent action of the indicated compounds on the clonal expansion of the MCF-7 cells expressing the wild-type ERα or the D538G of Y537S ER mutations (n = 3-8) in the absence **(A)** and presence **(B)** of 10 nM oestradiol.

**Supplementary Figure S6**

Effect of the AHR antagonist CH223191 on the inhibition of proliferation by the X compounds. Quantification of the action of the AHR antagonist CH223191 on the inhibition of the clonal expansion of MCF-7 cells by the indicated X compounds. Cells were treated with 10 nM of the X compounds in the absence and presence of 1 µM CH223191 for 14 days. The values are the means ± SEM, n = 4. ***p< 0.001; ns is not significant.

Supplementary Figure 7

Generation of CRISPR/Cas9 knockout AHR. **A**. Strategy for the generation of AHR knockout. In strategy 1, gRNA _805 was chosen to introduce a double-stranded DNA break immediately downstream of the *AHR* start codon so that repair by nonhomologous end joining will introduce indels to disrupt the protein reading frame. In strategy 2, gRNA_708 upstream and gRNA_805 downstream of the *AHR* start codon should cause two double-stranded DNA breaks in approx. 100 bp distance. Thus, repair by nonhomologous end joining will delete the start codon and block transcription. **A. and B.** Selection of AHR KO clones in MCF7 and T47D cells. Western blot with equal numbers of cell clones isolated from the AHR CRISPR Cas9 knockout studies in MCF-7 **(A)** and T47D **(B)** cells with different sgRNA. Anti-AHR and β-actin antibodies were used for the Western blots.

**Supplementary Figure 8**

2-phenyl-imidazo[1, 2α] pyridine-mediated inhibition of ER^+^ T47D breast cancer cell proliferation via AHR. **A**. Proliferation of empty vector transfected T47D cells and T47D AHR KO clones #7 and #30 cultured in complete media and counted on days 0 and 5. The starting number of cells was 1 x10^4^ per well. Data are the averages of three independent experiments ± SEM, normalized to day 0., ****p ≤ 0.0001. **B.** Quantification of the concentration dependent action of the indicated compounds on the clonal expansion of empty vector transfected T47D cells and AHR KO T47D clones #7 and #30. The results are shown as bar charts and they represent the means ± SEM, n = 3-4. * p< 0.05; ** p < 0.01; *** p < 0.001; **** < 0.0001; ns is not significant.

## References

1. Bray, F. et al. Global cancer statistics 2022: GLOBOCAN estimates of incidence and mortality worldwide for 36 cancers in 185 countries. CA Cancer J Clin 74, 229–263 (2024).

2. Schettini, F. et al. Endocrine-Based Treatments in Clinically-Relevant Subgroups of Hormone Receptor-Positive/HER2-Negative Metastatic Breast Cancer: Systematic Review and Meta-Analysis. Cancers (Basel) 13, 1458 (2021).

3. Downton, T., Zhou, F., Segara, D., Jeselsohn, R. & Lim, E. Oral Selective Estrogen Receptor Degraders (SERDs) in Breast Cancer: Advances, Challenges, and Current Status. Drug Des Devel Ther 16, 2933–2948 (2022).

4. Hanker, A. B., Sudhan, D. R. & Arteaga, C. L. Overcoming Endocrine Resistance in Breast Cancer. Cancer Cell 37, 496–513 (2020).

5. Robinson, D. R. et al. Activating ESR1 mutations in hormone-resistant metastatic breast cancer. Nat. Genet. 45, 1446–1451 (2013).

6. Jeselsohn, R. et al. Emergence of constitutively active estrogen receptor-α mutations in pretreated advanced estrogen receptor-positive breast cancer. Clin Cancer Res 20, 1757–1767 (2014).

7. Toy, W. et al. Activating ESR1 Mutations Differentially Affect the Efficacy of ER Antagonists. Caner Discov 7, 277–287 (2017).

8. Rinaldi, J. et al. The genomic landscape of metastatic breast cancer: Insights from 11,000 tumors. PLoS One 15, e0231999 (2020).

9. Jeselsohn, R. et al. Allele-Specific Chromatin Recruitment and Therapeutic Vulnerabilities of ESR1 Activating Mutations. Cancer Cell 33, 173–186 (2018).

10. O’Leary, B. et al. The Genetic Landscape and Clonal Evolution of Breast Cancer Resistance to Palbociclib plus Fulvestrant in the PALOMA-3 Trial. Cancer Discov 8, 1390–1403 (2018).

11. Wang, Y. & Tang, S. C. The race to develop oral SERDs and other novel estrogen receptor inhibitors: recent clinical trial results and impact on treatment options. Cancer Metastasis Rev 41, 975–990 (2022).

12. Garcia-Fructuoso, I., Gomez-Bravo, R. & Schettini, F. Integrating new oral selective oestrogen receptor degraders in the breast cancer treatment. Curr Opin Oncol 34, 635– 642 (2022).

13. Hoy, S. M. Elacestrant: First Approval. Drugs 83, 555–561 (2023).

14. Oliveira, M. et al. Camizestrant, a next-generation oral SERD, versus fulvestrant in post-menopausal women with oestrogen receptor-positive, HER2-negative advanced breast cancer (SERENA-2): a multi-dose, open-label, randomised, phase 2 trial. Lancet Oncol 25, 1424–1439 (2024).

15. Tolaney, S. M. et al. AMEERA-3: Randomized Phase II Study of Amcenestrant (Oral Selective Estrogen Receptor Degrader) Versus Standard Endocrine Monotherapy in Estrogen Receptor-Positive, Human Epidermal Growth Factor Receptor 2-Negative Advanced Breast Cancer. J Clin Oncol 41, 4014–4024 (2023).

16. Martín, M. et al. Giredestrant for Estrogen Receptor-Positive, HER2-Negative, Previously Treated Advanced Breast Cancer: Results From the Randomized, Phase II acelERA Breast Cancer Study. J Clin Oncol 42, 2149–2160 (2024).

17. Pan, M. et al. Identification of an Imidazopyridine-based Compound as an Oral Selective Estrogen Receptor Degrader for Breast Cancer Therapy. Cancer Res Commun 3, 1378–1396 (2023).

18. Harrod, A. et al. Genome engineering for estrogen receptor mutations reveals differential responses to anti-estrogens and new prognostic gene signatures for breast cancer. Oncogene 41, 4905–4915 (2022).

19. Segatto, I. et al. A comprehensive luminal breast cancer patient-derived xenografts (PDX) library to capture tumor heterogeneity and explore the mechanisms of resistance to CDK4/6 inhibitors. J Pathol 264, 434–447 (2024).

20. Gruszczyk, J. et al. Cryo-EM structure of the agonist-bound Hsp90-XAP2-AHR cytosolic complex. Nat Commun 13, 7010 (2022).

21. Concordet, J. P. & Haeussler, M. CRISPOR: intuitive guide selection for CRISPR/Cas9 genome editing experiments and screens. Nucleic Acids Res 46(W1), W242–W245 (2018).

22. Cong, L. et al. Multiplex genome engineering using CRISPR/Cas systems. Science. 339, 819–823 (2013).

23. Guzmán, C., Bagga, M., Kaur, A., Westermarck, J. & Abankwa, D. ColonyArea: an ImageJ plugin to automatically quantify colony formation in clonogenic assays. PLoS One 9, e92444 (2014).

24. Mootha, V. K. et al. PGC-1alpha-responsive genes involved in oxidative phosphorylation are coordinately downregulated in human diabetes. Nat Genet 34, 267–273 (2003).

25. Subramanian, A. et al. Gene set enrichment analysis: a knowledge-based approach for interpreting genome-wide expression profiles. Proc Natl Acad Sci U S A 102, 15545– 15550 (2005).

26. Kwong, H. S. et al. Structural Insights into the Activation of Human Aryl Hydrocarbon Receptor by the Environmental Contaminant Benzo[a]pyrene and Structurally Related Compounds. J Mol Biol 436, 168411 (2024).

27. Wang, S. et al. A 155-plex high-throughput in vitro coregulator binding assay for (anti-)estrogenicity testing evaluated with 23 reference compounds. ALTEX 30, 145– 157 (2013).

28. Houtman, R. et al. Serine-305 phosphorylation modulates estrogen receptor alpha binding to a coregulator peptide array, with potential application in predicting responses to tamoxifen. Mol Cancer Ther 11, 805–816 (2012).

29. Sastry, G. M., Adzhigirey, M., Day, T., Annabhimoju, R. & Sherman, W. Protein and ligand preparation: parameters, protocols, and influence on virtual screening enrichments. J Comput Aided Mol Des 27, 221–234 (2013).

30. Søndergaard, C. R., Olsson, M. H., Rostkowski, M. & Jensen, J. H. Improved Treatment of Ligands and Coupling Effects in Empirical Calculation and Rationalization of pKa Values. J Chem Theory Comput 7, 2284–2295 (2011).

31. Friesner, R. A. et al. Extra precision glide: docking and scoring incorporating a model of hydrophobic enclosure for protein-ligand complexes. J Med Chem 49, 6177–6196 (2006).

32. Simon, C. G. J. et al. Mechanism of action, potency and efficacy: considerations for cell therapies. J Transl Med 22, 416 (2024).

33. Depypere, H. T. et al. The serum estradiol concentration is the main determinant of the estradiol concentration in normal breast tissue. Maturitas 81, 42–45 (2015).

34. Guan, J. et al. Therapeutic Ligands Antagonize Estrogen Receptor Function by Impairing Its Mobility. Cell 178, 949–963 (2019).

35. Ramadoss, P., Marcus, C. & Perdew, G. H. Role of the aryl hydrocarbon receptor in drug metabolism. Expert Opin Drug Metab Toxicol. 1, 9–21 (2005).

36. Zheng, J. & Lafontant-Joseph, O. IUPAC Digitized pKa Dataset, v2.2. Int. Union Pure Appl. Chem. (2024).

37. Diao, X. et al. Structural basis for the ligand-dependent activation of heterodimeric AHR-ARNT complex. Nat Commun 16, 1282 (2025).

38. Zhao, B., Degroot, D. E., Hayashi, A., He, G. & Denison, M. S. CH223191 is a ligand-selective antagonist of the Ah (Dioxin) receptor. Toxicol Sci 117, 393–403 (2010).

39. Meng, L. et al. Epoxomicin, a potent and selective proteasome inhibitor, exhibits in vivo antiinflammatory activity. Proc Natl Acad Sci U S A 96, 10403–10408 (1999).

40. Wang, E., Xiang, K., Zhang, Y. & Wang, X. Patient-derived organoids (PDOs) and PDO-derived xenografts (PDOXs): New opportunities in establishing faithful pre-clinical cancer models. J Natl Cancer Cent 2, 263–276 (2022).

41. Opitz, C. A., Holfelder, P., Prentzell, M. T. & Trump, S. The complex biology of aryl hydrocarbon receptor activation in cancer and beyond. Biochem Pharmacol 216, 115798 (2023).

42. Luecke-Johansson, S. et al. A Molecular Mechanism To Switch the Aryl Hydrocarbon Receptor from a Transcription Factor to an E3 Ubiquitin Ligase. Mol Cell Biol 37, e00630–16 (2017).

43. Chen, Z. et al. Carbidopa suppresses estrogen receptor-positive breast cancer via AhR-mediated proteasomal degradation of ERα. Invest New Drugs 40, 1216–1230 (2022).

44. Wormke, M. et al. The aryl hydrocarbon receptor mediates degradation of estrogen receptor alpha through activation of proteasomes. Mol Cell Biol 23, 1843–1855 (2003).

45. Wormke, M., Stoner, M., Saville, B. & Safe, S. Crosstalk between estrogen receptor alpha and the aryl hydrocarbon receptor in breast cancer cells involves unidirectional activation of proteasomes. FEBS Lett 478, 109–112 (2000).

46. Cano-Sánchez, J. et al. The Aryl Hydrocarbon Receptor Ligand 6-Formylindolo(3,2-b)carbazole Promotes Estrogen Receptor Alpha and c-Fos Protein Degradation and Inhibits MCF-7 Cell Proliferation and Migration. Pharmacology 108, 157–165 (2023).

47. Chen, C. et al. Aryl hydrocarbon receptor: An emerging player in breast cancer pathogenesis and its potential as a drug target. Mol Biol Rep 29, 11 (2024).

48. Safe, S. & Zhang, L. The Role of the Aryl Hydrocarbon Receptor (AhR) and Its Ligands in Breast Cancer. Cancers (Basel) 14, 5574 (2022).

49. Tiong, C. T. et al. A novel prenylflavone restricts breast cancer cell growth through AhR-mediated destabilization of ERα protein. Carcinogenesis 33, 1089–1097 (2012).

50. Cockroft, S. L., Hunter, C. A., Lawson, K. R., Perkins, J. & Urch, C. J. Electrostatic control of aromatic stacking interactions. J Am Chem Soc 127, 8594–8595 (2005).

51. Hunter, C. A., Lawson, K. R., Perkins, J. & Urch, C. J. Aromatic interactions. J. Chem. Soc. Perkin Trans. 2 651–669 (2001) doi:10.1039/B008495F.

52. Cozzi, F. et al. Through-space interactions between face-to-face, center-to-edge oriented arenes: importance of polar-pi effects. Org Biomol Chem 1, 157–162 (2003).

53. Chen, Z. et al. Carbidopa suppresses prostate cancer via aryl hydrocarbon receptor-mediated ubiquitination and degradation of androgen receptor. Oncogenesis 9, 49 (2020).

54. Zgarbová, E. & Vrzal, R. The Impact of Indoles Activating the Aryl Hydrocarbon Receptor on Androgen Receptor Activity in the 22Rv1 Prostate Cancer Cell Line. Int J Mol Sci 24, 502 (2022).

55. Sun, F. et al. A novel prostate cancer therapeutic strategy using icaritin-activated arylhydrocarbon-receptor to co-target androgen receptor and its splice variants. Carcinogenesis 36, 757–768 (2015).

56. Ohtake, F., Fujii-Kuriyama, Y. & Kato, S. AhR acts as an E3 ubiquitin ligase to modulate steroid receptor functions. Biochem Pharmacol 77, 474–484 (2009).

57. Ohtake, F. et al. Dioxin receptor is a ligand-dependent E3 ubiquitin ligase. Nature 446, 562–566 (2007).

58. Marconett, C. N. et al. Indole-3-carbinol triggers aryl hydrocarbon receptor-dependent estrogen receptor (ER)alpha protein degradation in breast cancer cells disrupting an ERalpha-GATA3 transcriptional cross-regulatory loop. Mol Biol Cell 21, 1166–1177 (2010).

59. Ohoka, N. et al. Development of Small Molecule Chimeras That Recruit AhR E3 Ligase to Target Proteins. ACS Chem Biol 14, 2822–2832 (2019).

60. Matthews, J. & Gustafsson, J. A. Estrogen receptor and aryl hydrocarbon receptor signaling pathways. Nucl Recept Signal e016, (2006).

