## Supplementary Tables and Figures for "Distinct 2-phenyl-imidazo[1, 2α] pyridine derivatives drive ER degradation and selectively impair proliferation of ER^+^ breast cancer cells via the aryl hydrocarbon receptor": Supplementary Figures .pdf

A

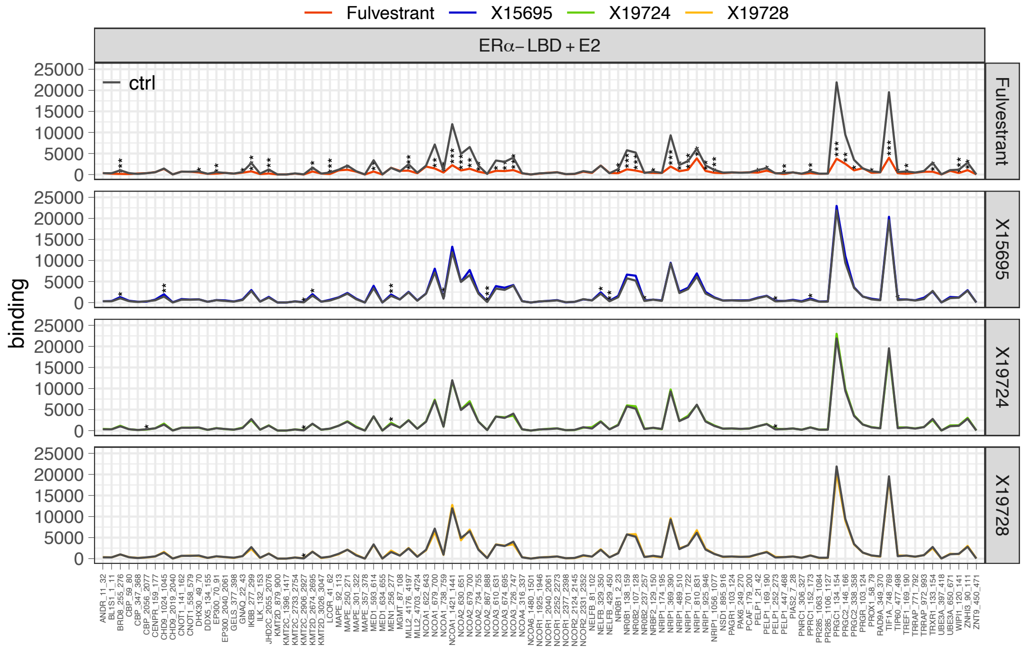


B

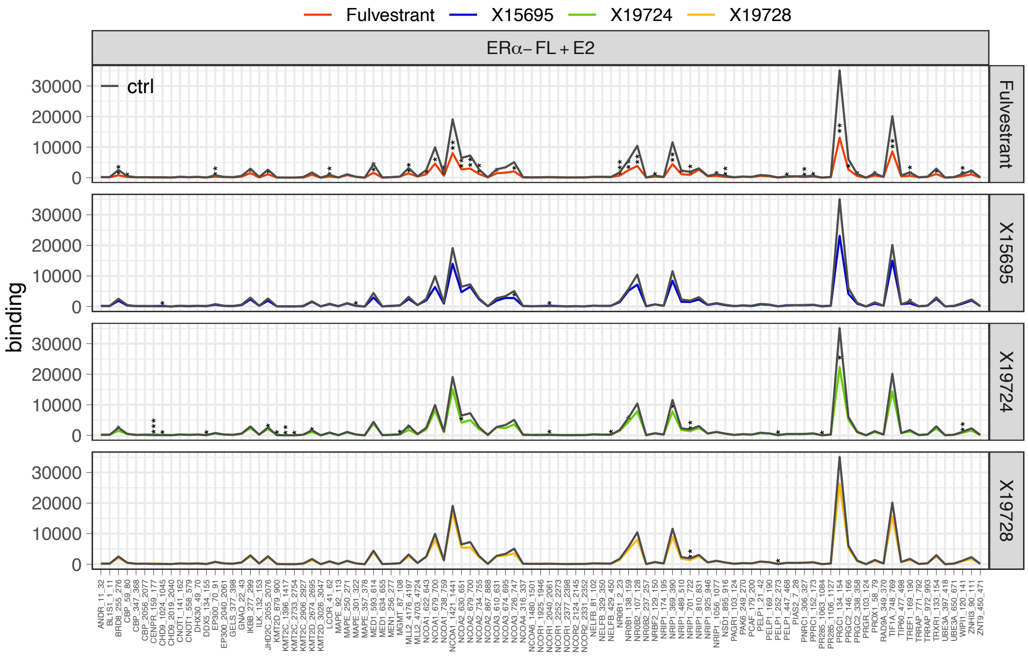


Supplementary 1

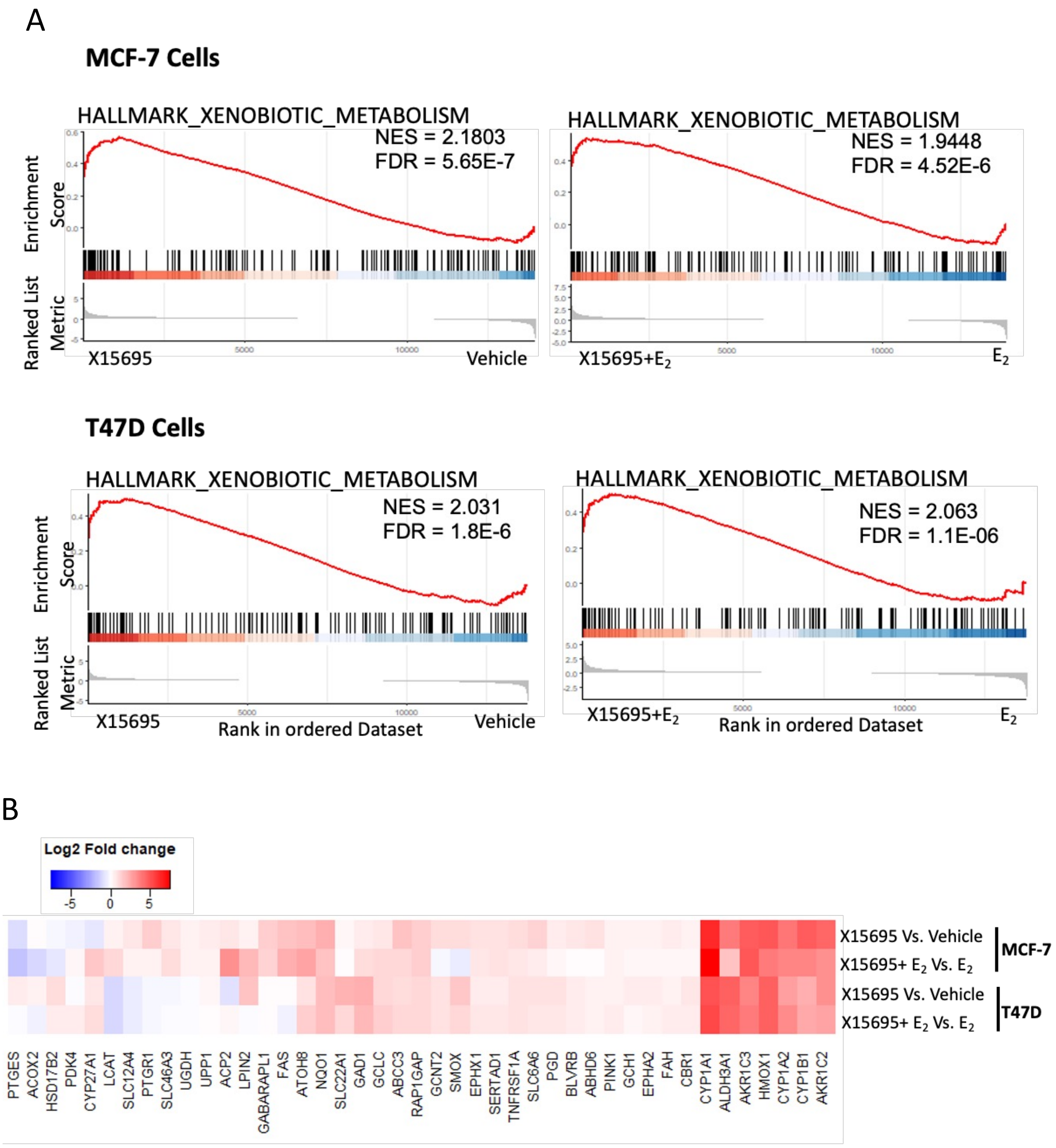


Supplementary Fig. 2

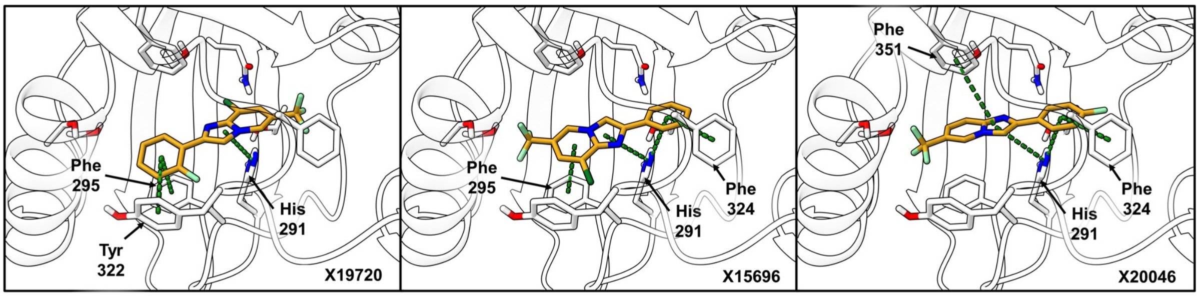


Supplementary Fig. 3

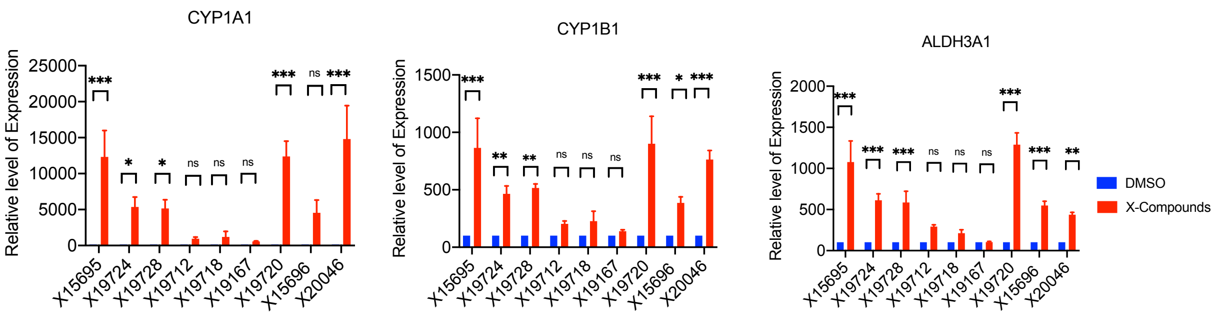


Supplementary Fig. 4

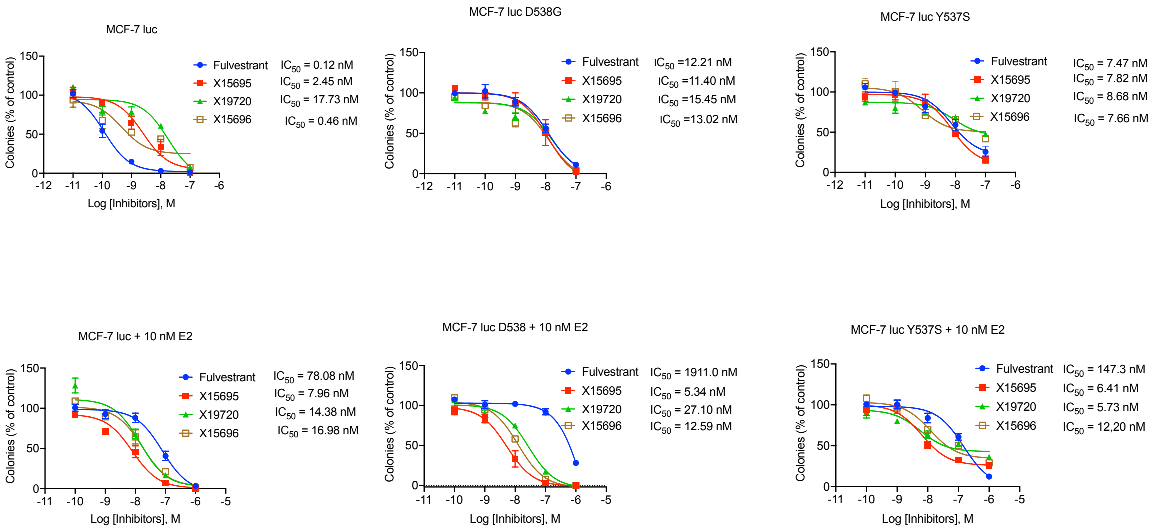


**Supplementary Fig. 5**

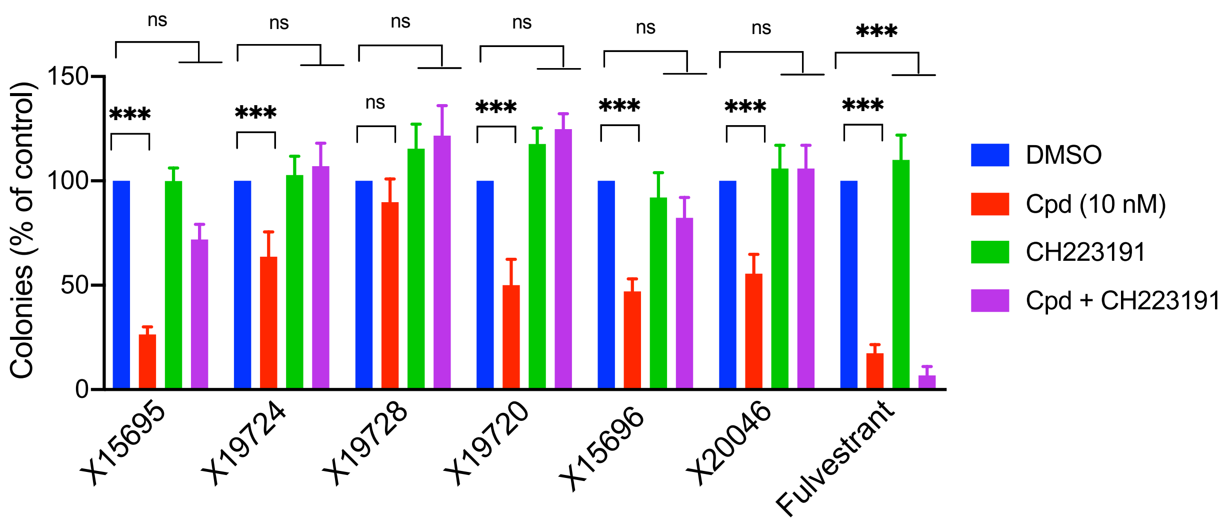


**Supplementary Figure 6**

A

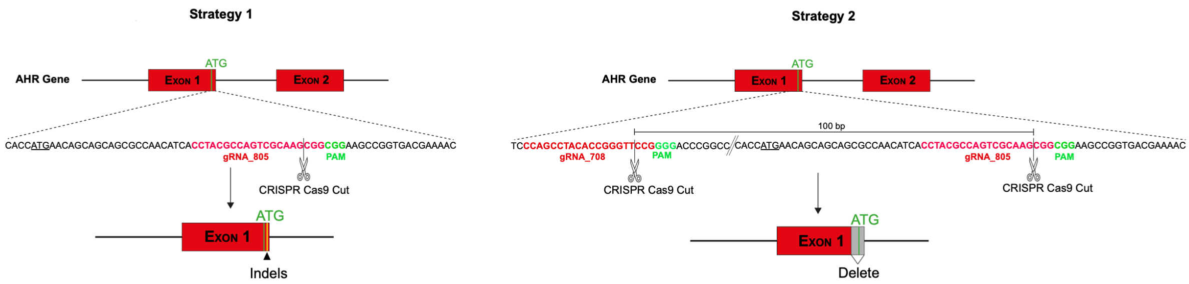


MCF-7 cells

B

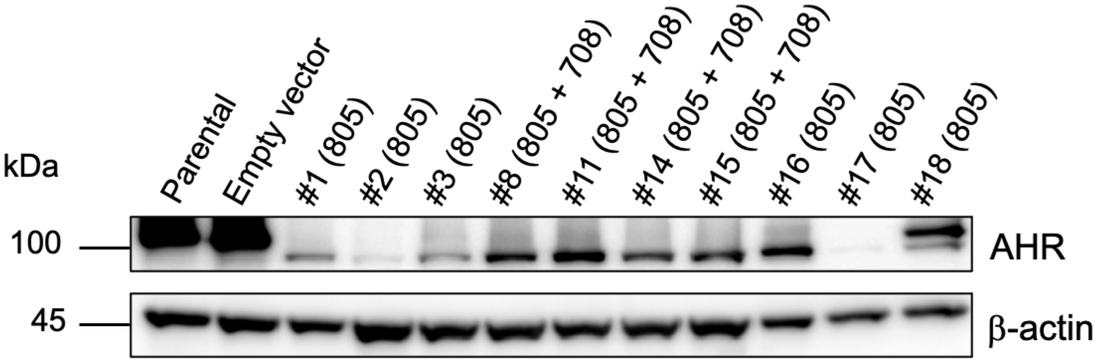


T47D cells

C

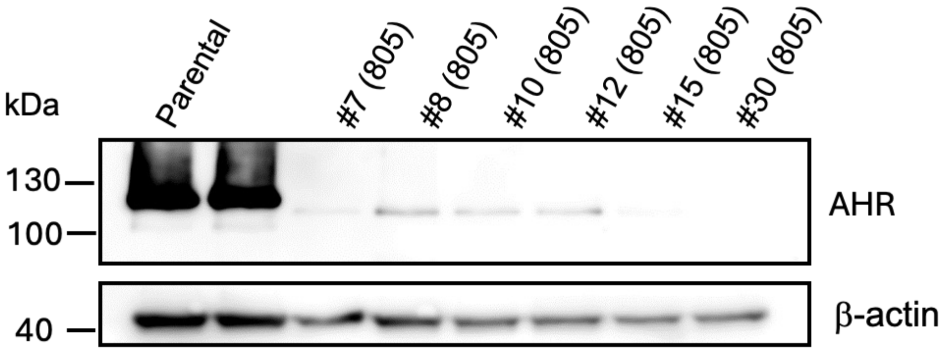


**Supplementary Figure 7**

A

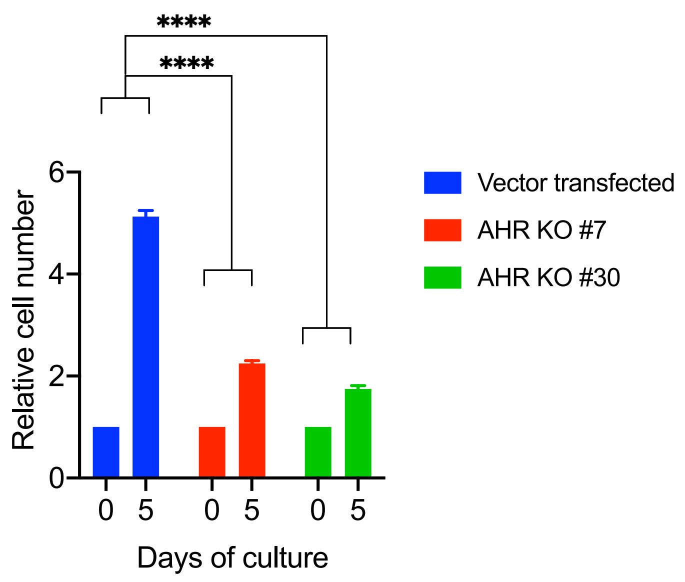


B

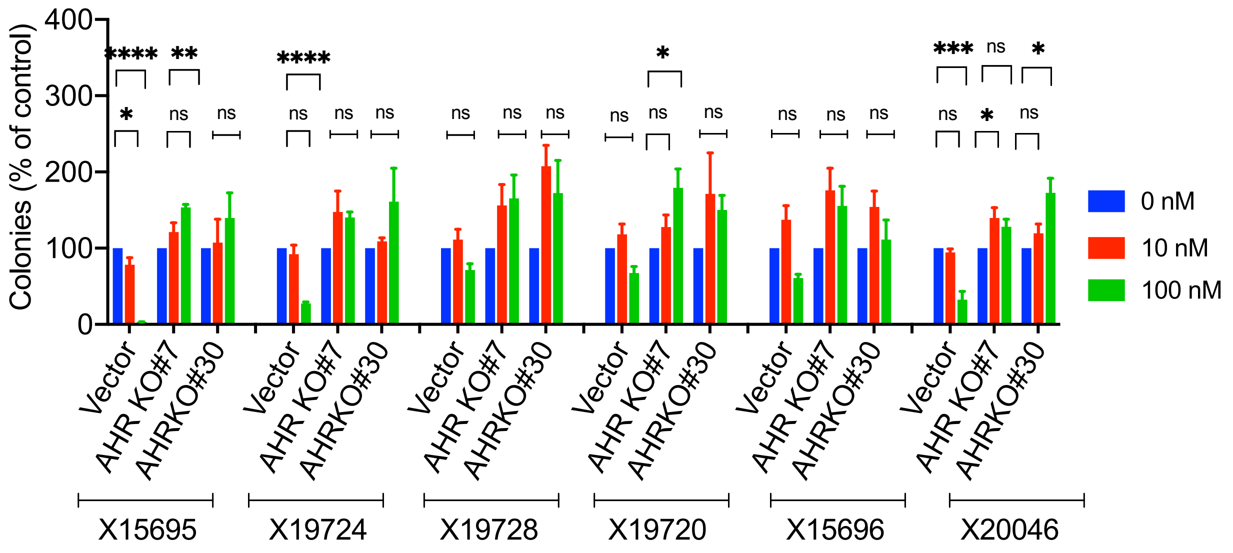


**Supplementary Figure 8**
