## Supplementary Tables and Figures for "Distinct 2-phenyl-imidazo[1, 2α] pyridine derivatives drive ER degradation and selectively impair proliferation of ER^+^ breast cancer cells via the aryl hydrocarbon receptor": Supplementary Tables.pdf

**Supplementary Table 1** Primers for ER and AhR target gene expression

| | |
|---|---|
| human PGR For<br>human PGR Rev | 5´-CTTAATCAACTAGGCGAGAG-3´<br>5´-AAGCTCATCCAAGAATACTG-3´ |
| human GREB For<br>human GREB Rev | 5´- GAGTGACAATGAGGAAGAG-3´<br>5´-CTCGTTGGAAATGGAGACAA-3´ |
| human PDZK1 For<br>human PDZK1 Rev | 5´-GCAGGCTCAGAACAGAAAGG-3´<br>5´- TCCAGGGTTTCCACAGACTC-3´ |
| human TFF1 For<br>human TFF1 Rev | 5´-CAATGGCCACCATGGAGAAC-3´<br>5´-AACGGTGTCGTCGAAACAGC-3´ |
| human Cyp1A1 For<br>human Cyp1A1 Rev | 5' TCAGCTCAGTACCTCAGCCA 3'<br>5' CATGGCCCTGGTGGATTCTT 3' |
| human CYP1B1 For<br>human CYP1B1 Rev | 5'CCACTATCACTGACATCTTC 3'<br>5' ACGACCTGATCCAATTCT 3' |
| human ALDH3A1 For<br>human ALDH3A1 Rev | 5' TGACTACATCCTCTGTGA 3'<br>5' GCACTAATGATTCTTCCATAG 3' |

**Supplementary Table 2**: Results of the linear fit for the calibration of X15695, X15696 and X19720.

| | **X15695** | **X15696** | **X19720** |
|---|---|---|---|
| **Best-fit values** | | | |
| Slope | 140.6 ± 9.248 | 146.4 ± 2.245 | 142.7 ± 2.497 |
| Y-intercept when X=0.0 | 3.910 ± 100.9 | 29.54 ± 24.49 | -25.01 ± 27.24 |
| X-intercept when Y=0.0 | -0,02781 | -0,2018 | 0,1753 |
| 1/slope | 0,007112 | 0,006831 | 0,007007 |
| **95% Confidence Intervals** | | | |
| Slope | 120.4 to 160.8 | 141.5 to 151.3 | 137.3 to 148.1 |

| | | | |
|---|---|---|---|
| Y-intercept when X=0.0 | -216.0 to 223.8 | -23.82 to 82.90 | -84.36 to 34.34 |
| X-intercept when Y=0.0 | -1.767 to 1.413 | -0.5792 to 0.1593 | -0.2468 to 0.5772 |
| **Goodness of Fit** | | | |
| R square | 0,9506 | 0,9972 | 0,9963 |
| Sy.x | 287.1 | 69.67 | 77.49 |
| **Slope significantly non-zero?** | | | |
| F | 231,1 | 4253 | 3267 |
| DFn, DFd | 1.000, 12.00 | 1.000, 12.00 | 1.000, 12.00 |
| P value | < 0.0001 | < 0.0001 | < 0.0001 |
| Deviation from zero? | Significant | Significant | Significant |
| **Data** | | | |
| Number of X values | 7 | 7 | 7 |
| Maximum number of Y replicates | 2 | 2 | 2 |
| Total number of values | 14 | 14 | 14 |
| Number of missing values | 4 | 4 | 4 |

**Supplementary Table 3**: Results of measurements (N = 2) on the thermodynamic solubility of analytes X15695, X15696, and X19720. The retention time of each analyte (Rt), the corresponding mass-to-charge ratio (m/z), and the Area Under the Curve (AUC) of the chromatographic peak were determined.

| | **N = 1** | | | **N = 2** | | | **Mean** |
|---|---|---|---|---|---|---|---|
| | **Rt / min** | **AUC** | **m/z found** | **Rt / min** | **AUC** | **m/z found** | **AUC** |
| **X15695** | 5.968 | 532.606 | 314.8 | 5.996 | 391.862 | 314.7 | 423.653 |
| **X15696** | 5.927 | 783.832 | 297.1 | 5.930 | 771.367 | 297.0 | 777.600 |
| **X19720** | 6.188 | 164.348 | 315.2 | 6.192 | 172.102 | 315.0 | 168.225 |

**Supplementary Table 4**: Measured values of the stability test for the compounds X15695, X15696 and X19720. The retention times (Rt) of the substance peaks and the area under the curve (AUC) of the corresponding peak are shown.

| **Time / h** | **X15695** | | **X15696** | | **X19720** | |
|---|---|---|---|---|---|---|
| | **Rt / min** | **AUC** | **Rt / min** | **AUC** | **Rt / min** | **AUC** |
| **0** | 5.999 | 2495.299 | 5.920 | 1478.322 | 6.190 | 1234.075 |
| **24** | 6.002 | 2502.122 | 5.923 | 1484.881 | 6.198 | 1245.881 |
| **48** | 5.993 | 2505.480 | 5.911 | 1483.666 | 6.186 | 1248.872 |
| **72** | 5.991 | 2509.509 | 5.912 | 1486.572 | 6.182 | 1247.594 |
| **96** | 5.994 | 2510.456 | 5.913 | 1479.954 | 6.181 | 1243.603 |
| **120** | 5.993 | 2514.400 | 5.913 | 1485.339 | 6.178 | 1245.513 |
